# RNF213 M1-ubiquitinates damaged mitochondria to activate pro-inflammatory NF-κB signaling

**DOI:** 10.64898/2026.09.21.752888

**Authors:** Rosalie Heilig, Hannah Glover, Charlotte E L Mellor, Catherine Cloix, Michael J Mcilwraith, Lyndsey Flanagan, Nikki Paul, Peter Thomason, Bodhayan Prasad, Asma Ahmed, Hugo Dupuis, Destiny Dalseno, Leah Smith, James Hamilton, Ella Hall-Younger, Esmee Vringer, Lori Buetow, Alba Roca-Portoles, Alfredo Montes Gomes, Graeme Clark, Annabel Black, Sergio Lilla, Sara Zanivan, Leo Carlin, Vignir Helgason, Danny Huang, Stephen WG Tait

## Abstract

Through mitochondrial outer membrane permeabilization (MOMP), mitochondria are central to apoptosis. MOMP is pro-inflammatory, impacting diverse processes including anti-tumor immunity and cellular senescence. Despite this, how damaged mitochondria activate inflammation remains unclear. Recent studies have shown that following MOMP, mitochondria are extensively ubiquitinated activating pro-inflammatory NF-κB signaling. We identify the E3 ligase, RNF213, as essential for pro-inflammatory mitochondrial ubiquitination following MOMP. RNF213 has an established role in pro-inflammatory, cell-autonomous immunity to bacteria and other pathogens. We discover that RNF213 is required for mitochondrial M1-linked (linear) ubiquitination. Interestingly, this was found to be independent of the canonical M1-ubiquitin ligase complex LUBAC. RNF213 can directly catalyze M1-linked ubiquitination. By promoting mitochondrial M1-linked ubiquitination, RNF213 initiates the recruitment of the essential NF-κB adaptor molecule, NEMO, and subsequently activates pro-inflammatory NF-κB signaling. Collectively, our findings highlight striking similarities between how cells detect damaged mitochondria and intracellular pathogens, leading to inflammation.

## Introduction

Mitochondria are essential for the mitochondrial (intrinsic) pathway of apoptosis ^1^. During apoptosis, the pro-apoptotic BCL-2 family members BAX and BAK cause mitochondrial outer membrane permeabilization (MOMP), leading to cytochrome *c-*dependent caspase activation and cell death. Although apoptosis is considered an immunosilent cell death, paradoxically, MOMP itself is inherently pro-inflammatory ^2, 3^. Apoptotic caspase activity inhibits inflammation during cell death ^4–7^. Exploiting this, we and others find that killing cancer cells under caspase-inhibition promotes anti-tumor immunity dependent upon MOMP-induced inflammation in the dying tumor cell ^7–9^. MOMP-induced inflammation also has non-lethal biological functions in processes including senescence, immune surveillance and innate immunity ^10–12^. Nonetheless, how damaged mitochondria engage inflammation remains unclear.

Following MOMP, permeabilized mitochondria are extensively ubiquitinated ^13, 14^. Mitochondrial ubiquitination is promiscuous, affecting hundreds of mitochondrial proteins localized to both the inner and outer mitochondrial membranes. Mitochondrial ubiquitination recruits the ubiquitin-binding NF-κB adaptor molecule, NEMO (also called IKKγ), driving pro-inflammatory NF-κB signaling ^13^. NEMO has been shown to be activated by its M1-linked ubiquitination and by binding to M1-linked ubiquitin chains via its UBAN domain ^15^, in what is a critical step of TNF-dependent NF-κB signaling ^16, 17^. M1-linked ubiquitination is an atypical chain structure formed by linking ubiquitin molecules via the C-terminal carboxyl group of one ubiquitin, and the N-terminal methionine of the next ^18^. This specific ubiquitin chain linkage is generated by HOIP (HOIL-1 interacting protein, also known as RNF31), which is the catalytic subunit of the linear ubiquitin chain assembly complex (LUBAC, also comprised of HOIL-1 and SHARPIN)-the canonical E3 ligase complex for M1-linked ubiquitination ^17, 19^. Our previous study identified an enrichment of both K63 and M1-linked (linear) ubiquitin at the mitochondria following MOMP ^13^. We showed that a NEMO mutant which disrupts the UBAN domain and M1-ubiquitin binding (D311N) failed to be recruited to mitochondria following MOMP, indicating a role for M1-linked ubiquitin. However, NEMO recruitment was not impaired in HOIP-deficient MEF cells, arguing against a role for LUBAC in this process ^13^. Further investigations revealed that MOMP-dependent mitochondrial ubiquitination was also independent of the PINK1-Parkin pathway and mitochondrial resident E3 ligases MUL1 (also called MAPL) and MARCH5 ^13^. Therefore, the E3 ligase machinery responsible for MOMP-dependent ubiquitination of mitochondria remained elusive.

In this study, we set out to further investigate, define and characterize the E3 ubiquitin ligase machinery required for mitochondrial ubiquitination following MOMP. We identify RNF213 - a giant E3 ligase - as associating specifically with permeabilized mitochondria. Amongst other roles, RNF213 functions in pro-inflammatory, cell-autonomous immunity against diverse pathogens including bacteria, viruses and parasites ^20–23^. We find that RNF213 is essential for the M1-linked ubiquitination which enables mitochondrial NEMO recruitment upon MOMP. Surprisingly, we find that this M1 mitochondrial ubiquitination requires RNF213 but is independent of LUBAC. RNF213 can therefore function as an M1-ubiquitin ligase upon mitochondrial permeabilisation. We show that loss of RNF213 impairs both MOMP-induced NF-κB activity and subsequent inflammation. Possibly relating to the bacterial ancestry of mitochondria, our data reveal unexpected similarities between how damaged mitochondria and intracellular pathogens elicit inflammation.

## Results

### RNF213 ubiquitin ligase associates with and ubiquitinates permeabilized mitochondria upon MOMP

Upon MOMP, mitochondria are extensively ubiquitinated leading to NEMO recruitment and pro-inflammatory NF-κB signaling, as we have previously found ^13^. This ubiquitination was independent of the PINK1-Parkin pathway, cIAP1/2, XIAP and mitochondrial resident E3 ubiquitin ligases MUL1 (also called MAPL) and MARCH5 ^13^. From this, we reasoned that a relevant E3 ligase may be specifically recruited to permeabilized mitochondria. To examine this, we used MITO-tag (mitochondrial localized 3xHA epitope tag) ^24^ to define the mitochondrial proteome from cells undergoing MOMP, following treatment with a combination of BH3-mimetics, ABT-737 (ABT; inhibits BCL-xL, BCL-2, BCL-w) and S63845 (S6; inhibits MCL-1) in the presence of caspase inhibitor Q-VD-OPh (QVD) (**Fig. 1A**). SVEC4-10 mouse endothelial cells (hereafter called SVEC) stably expressing MITO-tag were generated; anti-HA immunoprecipitation enabled effective isolation of mitochondria, demonstrated by western blot of specific mitochondrial proteins such as cytochrome *c*, VDAC1/2, CypD, CVa and Core1 (**Fig. 1B**). MITO-tag expressing SVEC cells were treated to undergo MOMP, and isolated mitochondria were then subjected to mass-spectrometry analysis (**Fig. 1C**). Verifying engagement of mitochondrial apoptosis, the pro-apoptotic BCL-2 family member BAX was enriched, whereas the intermembrane space proteins cytochrome *c*, SMAC/DIABLO, HTRA2/OMI and AK2 were lost from mitochondria isolated from BH3-mimetic treated cells (**Fig. 1C**). We next asked whether any E3 ubiquitin ligases were significantly enriched on mitochondria following MOMP. Various E3 ubiquitin ligases were found to associate with mitochondria following BH3-mimetic treatment (**Fig. 1C)**. To reduce the set of potential E3-ligase candidates we further analysed a dataset previously generated to identify ubiquitinated proteins following MOMP^13^, on the premise that ubiquitination may indicate E3 ligase activation. This revealed that of the E3 ligases found to associate with mitochondria, only RNF213 and TRIM56 became ubiquitinated upon MOMP (**Fig. 1D**). RNF213 (also called Mysterin) is the largest known human E3 ubiquitin ligase (∼600 kDa), comprising a large N-terminal disordered region, a dynein-like AAA+ core, and a C-terminal E3 module containing the RING and RZ domains. RNF213 is the major susceptibility gene for moyamoya disease, a cerebrovascular disorder characterized by progressive intracranial arterial stenosis and collateral vessel formation ^25^. With roles in lipotoxicity, lipid droplet formation, and inflammation, RNF213 also has well-established functions in innate immunity during pathogen infection ^23, 26–28^, prompting us to investigate its role in mitochondrial damage-induced signalling. To further investigate RNF213 localisation, MEF expressing GFP-RNF213 were treated with BH3-mimetics to engage MOMP, and immunostained with anti-TOM20 antibody to visualise mitochondria. Cells were imaged by Airyscan (**Fig. 1E**) or Elyra super resolution microscopy, followed by image analysis by Imaris including surface rendering, segmentation and surface overlap quantification (**Fig. 1F, G**). This revealed mitochondrial overlap of GFP-RNF213 specifically following MOMP, consistent with our mass spec. analysis (**Fig 1E – 1G**). These data demonstrate that the E3 ligase RNF213 associates with mitochondria and is ubiquitinated upon MOMP.

**Figure 1.**
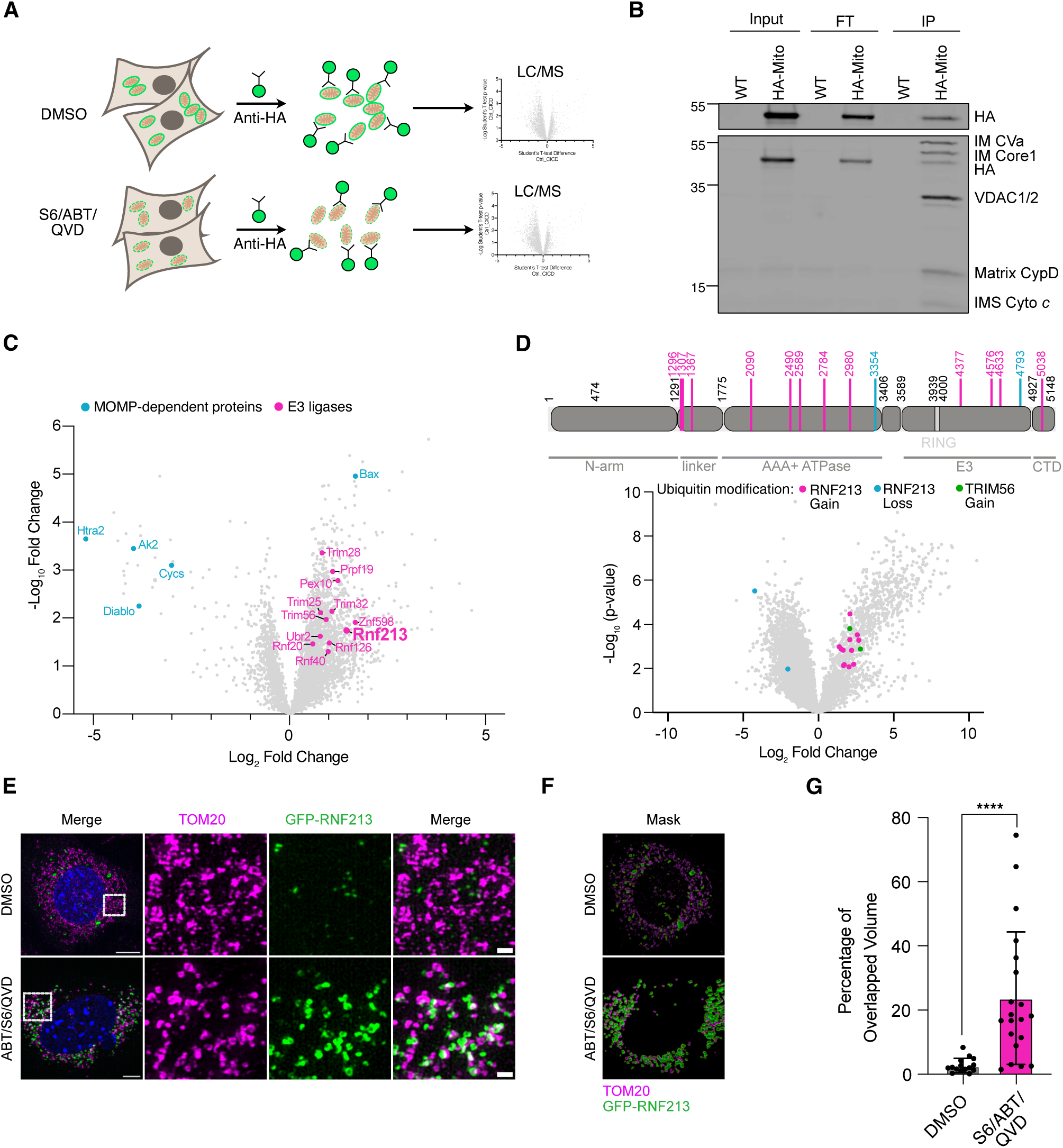
RNF213 is associated with the mitochondria and is ubiquitinated upon MOMP. **A)** Schematic illustrating rapid immunopurification of mitochondria using SVEC cells expressing 3xHA-EGFP-OMP25 upon treatment with DMSO or the combination of inhibitors S63845, ABT-737, Q-VD-OPh and subsequent proteomic analysis by LC/MS. **B)** Validation of immunopurification of mitochondria. SVEC WT or 3xHA-EGFP-OMP25 expressing cells were treated for 1 hr +/- ABT-737, S63845 and Q-VD-OPh and mitochondria were immunopurified with anti-HA beads. Lysates were immunoblotted for HA and mitochondrial proteins using membrane integrity antibody cocktail. **C)** Volcano plot of proteins in SVEC cells treated for 1 hr +/- ABT-737, S63845 and Q-VD-OPh. Proteins of interest are highlighted. The experiment was performed with n = 4 independent repeats. **D)** Schematic of murine RNF213 and volcano plot of ubiquitinated peptides in SVEC cells treated for 3 hrs +/- ABT-737, S63845 and Q-VD-OPh. Peptides containing amino acid residues of murine RNF213 with increased ubiquitination in DMSO (blue) or ABT-737, S63845, Q-VD-OPh (pink) treated cells are highlighted. Peptides containing amino acid resides of murine TRIM56 with increased ubiquitination in ABT-737, S63845, Q-VD-OPh (green) treated cells are highlighted. The experiment was performed with n = 4 independent repeats. **E)** MEF cells were induced to express GFP-RNF213 using 1 µg/ml doxycycline for >15 hrs. Cells were then treated +/- ABT-737, S63845 and Q-VD-OPh for 3 hrs and imaged by Elyra7 lattice structured illumination microscopy. GFP-RNF213 (green), TOM20 (magenta). Scale bar = 10 µm on main image, 1 µm on insets. **F)** Example Imaris rendering of cells imaged in E. GFP-RNF213 (green), TOM20 (magenta). **G)** Percentage of overlapped volume between GFP-RNF213 and TOM20, measured from Imaris rendering in F. Statistics were performed using one-way ANOVA. ****P<0.001.

### RNF213 promotes NF-κB dependent-inflammation following MOMP

Since RNF213 is recruited to mitochondria and ubiquitinated, we next asked whether it could promote MOMP-induced inflammation. RNF213-deleted SVEC cells were generated by CRISPR/Cas9 genome editing, hereafter called RNF213 knockout (KO). Efficient deletion of RNF213 was confirmed at the protein level by mass-spectrometry and at the genomic level by ICE analysis (**S1A – C**). Incucyte live-cell imaging demonstrated that RNF213 did not affect mitochondrial apoptosis following combined BH3-mimetic treatment (**Fig. S1D**). RNA-seq revealed no significantly differentially expressed genes between WT (wild type) and RNF213 KO SVEC cells under basal conditions (data not shown). Next, SVEC cells were treated to undergo MOMP and analyzed by RNA-seq (**Fig. 2A**). As expected, MOMP led to potent up-regulation of many pro-inflammatory transcripts and gene-ontology (GO) analysis highlighted GO-terms consistent with an anti-pathogen response (**Fig. 2B, Fig. S1E**). Importantly, RNF213 deletion blunted the transcriptional inflammatory response (**Fig. 2C**). We next asked which inflammatory pathways are promoted by RNF213 following MOMP. Control or RNF213 KO SVEC cells were analyzed for NF-κB activation by measuring phospho-IκBα (p-IκBα) by western blot (**Fig. 2D**). In RNF213 KO cells, p-IκBα levels were reduced, consistent with inhibition of NF-κB signaling. We and others have shown that following MOMP, subsequent release of mitochondrial DNA (mtDNA) activates cGAS-STING signaling ^2, 3, 5, 6^. cGAS-STING can also activate NF-κB ^29^. We therefore asked whether RNF213 regulates cGAS-STING activation following MOMP by measuring the level of phospho-STING (pSTING), phospho-TBK1 (pTBK1) and phospho-IRF3 (pIRF3), which are downstream outputs of cGAS-STING activity (**Fig. 2E**). RNF213 loss did not affect pSTING, pTBK1 or pIRF3 levels, indicating that RNF213 does not impact mtDNA-dependent cGAS-STING activation. To confirm that RNF213 contributes to the NF-κB-dependent inflammatory output, WT and RNF213 KO SVEC cells were treated to undergo MOMP. We have previously defined NF-κB and cGAS-STING dependent inflammatory transcripts following MOMP ^7^. Using this gene-set, the mRNA transcript and protein level of pro-inflammatory cytokines were measured by qRT-PCR (**Fig. 2F, Fig. S1F**) and multiplex cytokine assay (**Fig. 2G)**. Importantly, the levels of NF-κB regulated cytokines were reduced following MOMP at both transcript level (*Mcp1*, *Csf2*, *Cxcl1*) and protein level (MCP-1, GM-CSF, KC) in RNF213 KO cells. In contrast, the cGAS-STING regulated cytokines were unaffected at both transcript level (*Mip1a*, *Cxcl10*) and respective protein level (MIP-1-α, IP-10). Therefore, upon MOMP, RNF213 specifically promotes inflammatory NF-κB signaling.

**Figure 2.**
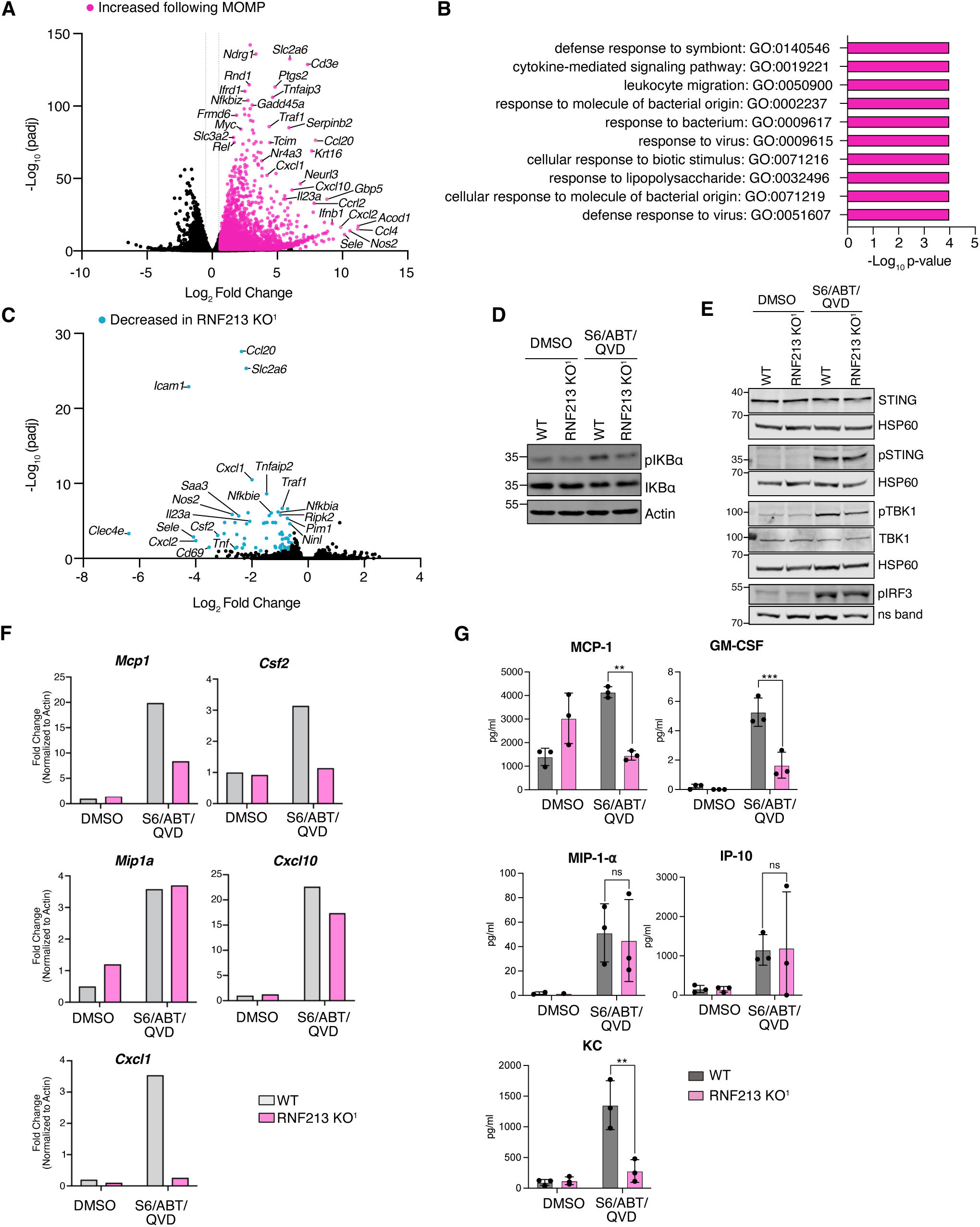
RNF213 promotes MOMP induced NF-κB activation. **A)** SVEC cells were treated for 3 hrs +/- ABT-737, S63845 and Q-VD-OPh. Bulk RNA was isolated and subjected to RNAseq analysis. Transcripts enriched following treatment are shown in pink. Data are averaged from 4 independent repeats. **B)** Gene-ontology analysis of the top upregulated transcripts (p adj < 10e^-4^) following ABT-737, S63845 and Q-VD-OPh treatment of SVEC cells. **C)** SVEC WT or RNF213 KO^1^ cells were treated for 3 hrs +/- ABT-737, S63845 and Q-VD-OPh. Bulk RNA was isolated and subjected to RNAseq analysis. Transcripts decreased following treatment in RNF213 KO^1^ cells are shown in blue. Data are averaged from 4 independent repeats. **D)** SVEC WT or RNF213 KO^1^ cells were treated for 1 hr +/- S63845, ABT-737 and Q-VD-OPh then immunoblotted for indicated antibodies. **E)** SVEC WT or RNF213 KO^1^ cells treated for 1 hr +/- S63845, ABT-737 and Q-VD-OPh then immunoblotted for indicated antibodies. The pIRF3 antibody gives a non-specific (ns) band, used as the loading control for this antibody. **F)** SVEC WT or RNF213 KO^1^ cells were treated for 3 hrs +/- ABT-737, S63845 and Q-VD-OPh. Expression of denoted transcripts was measured by qRT-PCR, normalised to actin. Graphs are representative of three independent experiments (related to figure S1D). **G)** SVEC WT or RNF213 KO^1^ cells treated for 8 hrs +/- S63845, ABT-737 and Q-VD-OPh. The levels of MCP-1, GM-CSF, MIP1a, IP-10 and Kc were measured using multiplex ELISA analysis. Graphs display mean values ± SD of n = 3 independent experiments. Statistics were performed using two-way ANOVA **P<0.01, ***P<0.005.

### RNF213 is required for mitochondrial ubiquitination following MOMP

Our previous study revealed that following MOMP, NF-κB activation is promoted by mitochondrial ubiquitination ^13^. As RNF213 drives MOMP-driven NF-κB activity, we investigated the contribution of RNF213 to mitochondrial ubiquitination. WT and RNF213 KO SVEC cells were treated to undergo MOMP and cytosolic and mitochondrial enriched fractions were analyzed by western blot using antibodies recognizing total-ubiquitin and K63-linked ubiquitin. Extensive ubiquitination of mitochondria occurred during MOMP in WT cells, detectable by both total and K63-linkage specific antibodies (**Fig. 3A, 3B**). RNF213 deletion reduced K63-linked and total mitochondrial ubiquitination. We next investigated whether RNF213 promotes mitochondrial ubiquitination in another cell type, using CRISPR-genome editing to delete RNF213 in mouse prostate cancer (CP2) cells, confirming RNF213 deletion at the genomic and protein level (**Fig. S2A - C**) ^30^. CP2 cells were treated to undergo MOMP, then cytosolic and mitochondrial enriched fractions were immunoblotted for ubiquitin (**Fig. 3C**). RNF213 KO CP2 cells also showed reduced mitochondrial ubiquitination. These data demonstrate that RNF213 promotes mitochondrial ubiquitination following MOMP.

**Figure 3.**
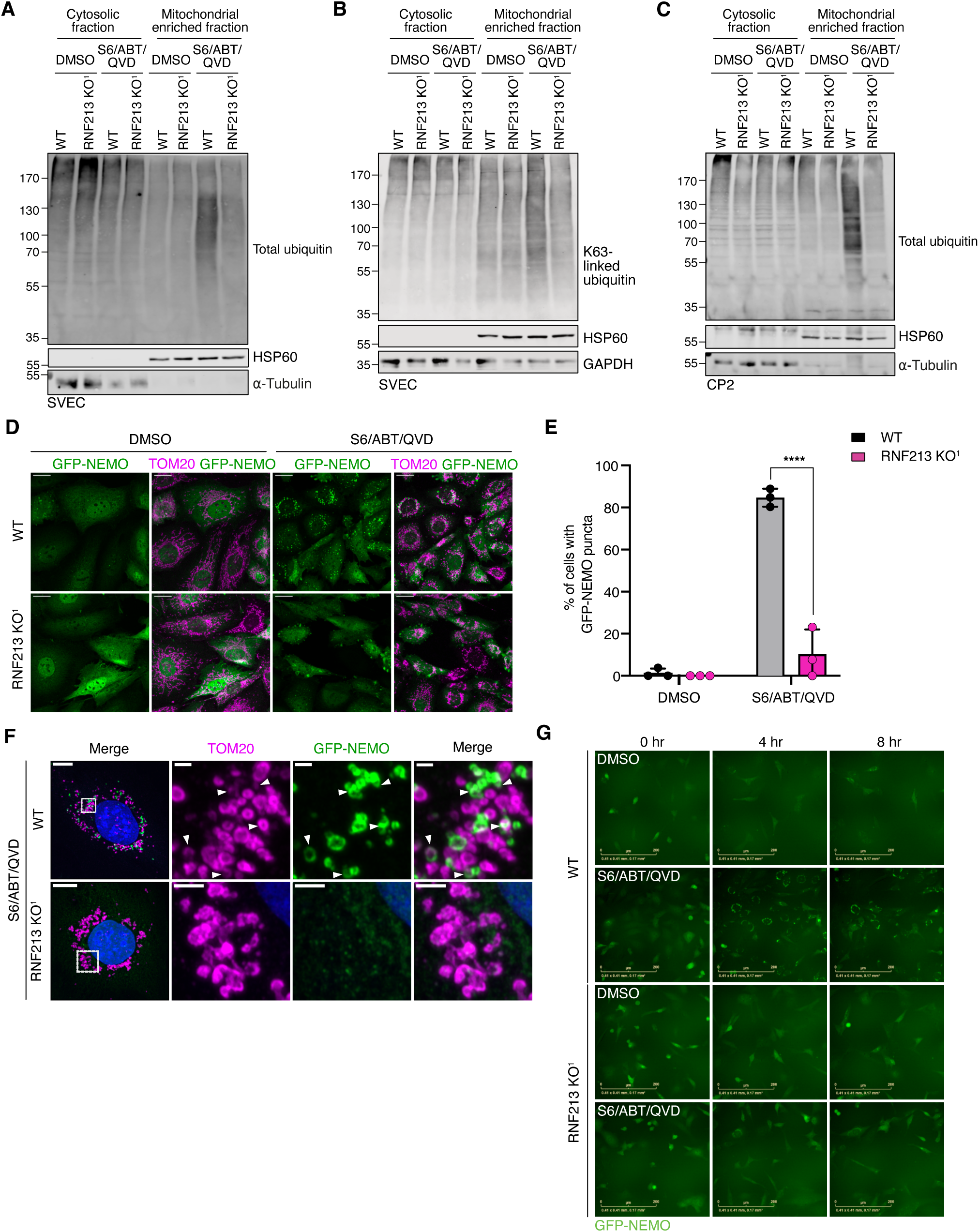
RNF213 promotes mitochondrial ubiquitination and is required for NEMO recruitment to mitochondria following MOMP. **A)** SVEC WT or RNF213 KO^1^ cells were treated +/- ABT-737, S63845 and Q-VD-OPh for 1 hr. Cytosolic/mitochondrial enriched fractions were obtained by Dounce fractionation and immunoblotted for total ubiquitin. **B)** SVEC WT or RNF213 KO^1^ cells were treated +/- ABT-737, S63845 and Q-VD-OPh for 1 hr. Cytosolic/mitochondrial enriched fractions were obtained by Dounce fractionation and immunoblotted for K63-linked ubiquitin. **C**) CP2 WT or RNF213 KO^1^ cells were treated +/- ABT-737, S63845 and Q-VD-OPh for 1 hr. Cytosolic/mitochondrial enriched fractions were obtained using digitonin fractionation and immunoblotted for total ubiquitin. Where indicated (A – C), HSP60, α-tubulin and GAPDH were probed as loading controls. **D)** SVEC WT or RNF213 KO^1^ cells expressing GFP-NEMO were treated +/- ABT-737, S63845 and Q-VD-OPh for 1 hr and imaged by LSM800 confocal microscopy. GFP-NEMO (green), TOM20 (magenta). Brightness and contrast of DMSO-treated images were independently adjusted from S6/ABT/QVD conditions to ensure optimal visualization of NEMO protein expression and mitochondrial localization. Scale bar = 10µm. **E)** GFP-NEMO mitochondrial puncta at mitochondria were quantified as described in methods using Image J/Fiji. SD from 3 independent experiments. Statistics were performed using two-way ANOVA: ****p<0.0001. **F)** SVEC WT or RNF213 KO^1^ cells expressing GFP-NEMO were treated with ABT-737, S63845 and Q-VD-OPh for 1 hr and imaged using LSM800 Airyscan super resolution microscopy. GFP-NEMO (green), TOM20 (magenta), DAPI (blue). Brightness and contrast of GFP-NEMO was increased in RNF213 KO^1^ cells to ensure optimal visualization of NEMO protein expression. Scale bar = 10µm, 1µm for insets. White arrows indicate TOM20 and GFP-NEMO overlap. **G)** SVEC WT or RNF213 KO^1^ cells expressing GFP-NEMO were treated +/- ABT-737, S63845 and Q-VD-OPh and imaged for 8 hrs using an Incucyte S3 to visualise mitochondrial NEMO recruitment. Indicated time points are shown. Scale bars = 200 µm. Experiments were carried out for a minimum of n = 3 independent repeats and representative images are shown.

### RNF213 is essential for mitochondrial NEMO association following MOMP

We previously found that NEMO recruitment to mitochondria requires ubiquitin binding ^13^. Given that RNF213 is important in the ubiquitination of permeabilized mitochondria, we asked if RNF213 was required for NEMO recruitment. RNF213 KO SVEC cells expressing GFP-NEMO were generated by CRISPR/Cas9 genome editing (**Fig. S2D**) and were treated to undergo MOMP. GFP-NEMO associated with mitochondria following MOMP, consistent with our previous data ^13^ (**Fig. 3D**). Crucially, RNF213 loss blocked mitochondrial GFP-NEMO recruitment. This was verified by quantifying GFP-NEMO puncta at mitochondria after 1hr (**Fig. 3E**). We next used Airyscan super-resolution imaging to image GFP-NEMO and mitochondria following MOMP in WT and RNF213-deleted SVEC cells (**Fig. 3F**). We observed GFP-NEMO puncta at mitochondria associating with the outer membrane protein TOM20 in an RNF213-dependent manner. To determine whether RNF213 is essential for sustained mitochondrial recruitment of NEMO, WT and RNF213 KO SVEC cells were induced to undergo MOMP and monitored by Incucyte live-cell image analysis over an extended period (**Fig. 3G**, **Movies S1 - S4**). No GFP-NEMO was observed at permeabilized mitochondria (denoted by punctate localization) in RNF213 KO SVEC cells, indicating that this recruitment is fully dependent on RNF213 (**Fig. 3G** and **Movies S2 and S4**). Thus, RNF213 is essential for mitochondrial ubiquitination and subsequent NEMO association following MOMP.

### RNF213 is required for mitochondrial M1-linked ubiquitination

NEMO has a much greater affinity for linear (M1) over K63-linked ubiquitin chains ^15^. We have previously reported that MOMP also causes M1-linked mitochondrial ubiquitination ^13^, therefore, we decided to confirm this observation and investigate it further. SVEC cells were treated to undergo MOMP and immunoblotted for M1-linked ubiquitin in cytosolic and mitochondrial enriched fractions, confirming that M1-linked ubiquitin is greatly increased (**Fig. 4A**). The specificity of the M1-antibody was validated by blotting recombinant di-ubiquitin of different linkages (**Fig. S3A**) and this result was also observed using an alternate M1-linkage specific antibody ^31^ (**Fig. S3B**). Moreover, ectopic expression of OTULIN, an M1-linkage specific deubiquitinating (DUB) enzyme ^32^, led to a decrease in mitochondrial associated M1-linked ubiquitination in SVEC and CP2 cells (**Fig. 4B, Fig. S3C**). We next investigated whether loss of mitochondrial integrity was sufficient to elicit M1-ubiquitination. To this end, we used raptinal, a compound which can cause MOMP independently of BAX and BAK ^33^. As expected, M1-ubiquitination elicited by BH3-mimetic treatment was completely dependent on BAX and BAK, however raptinal treatment effectively led to M1 ubiquitination even in the absence of BAX and BAK **(Fig. 4C)**. Therefore, loss of mitochondrial integrity is sufficient to engage M1 ubiquitination. Increased M1 ubiquitination could be observed following MOMP during apoptosis, and this was enhanced by caspase inhibition (**Fig 4D**.) To determine the extent of M1-ubiquitination relative to total ubiquitin linkage types following MOMP, we incubated mitochondrial enriched fractions with OTULIN, catalytically inactive (C129A) OTULIN or USP2, a linkage-independent DUB (**Figure 4E, 4F**). Specifically following MOMP, observed M1-ubiquitination could be efficiently removed by active OTULIN, but not the catalytically inactive C129A OTULIN mutant. Whereas USP2 effectively removed all ubiquitin, OTULIN had no discernible effects on total ubiquitination levels, demonstrating that M1-linked ubiquitin is a minor part of the total upregulation of mitochondrial ubiquitination following MOMP (**Figures 4E, 4F).** We next investigated whether RNF213 is required for M1-linked ubiquitination. WT or RNF213 KO SVEC cells were treated to undergo MOMP and analyzed by immunoblot for M1-linked ubiquitin (**Fig. 4G**). Compared to the increase observed in WT SVEC cells, mitochondrial M1-linked ubiquitination was effectively blocked in RNF213 KO cells (**Fig. 4G**). This result was confirmed using both a second, independent M1-specific antibody (**Fig. S3D**) and SVEC RNF213 KO cells generated using independent guide sequences (**Fig. S3E, S3F**). RNF213-dependent M1-linked ubiquitination was also replicated in multiple additional cell lines: CP2, MC38, NIH-3T3 and A549 (**Fig. 4H, Fig. S3G - S3L**). Interestingly, M1-linked ubiquitination was also found to be dependent on RNF213 following treatment with raptinal (**Fig. S3M**). These data therefore demonstrate that loss of mitochondrial integrity promotes RNF213-dependent M1-linked ubiquitination.

**Figure 4.**
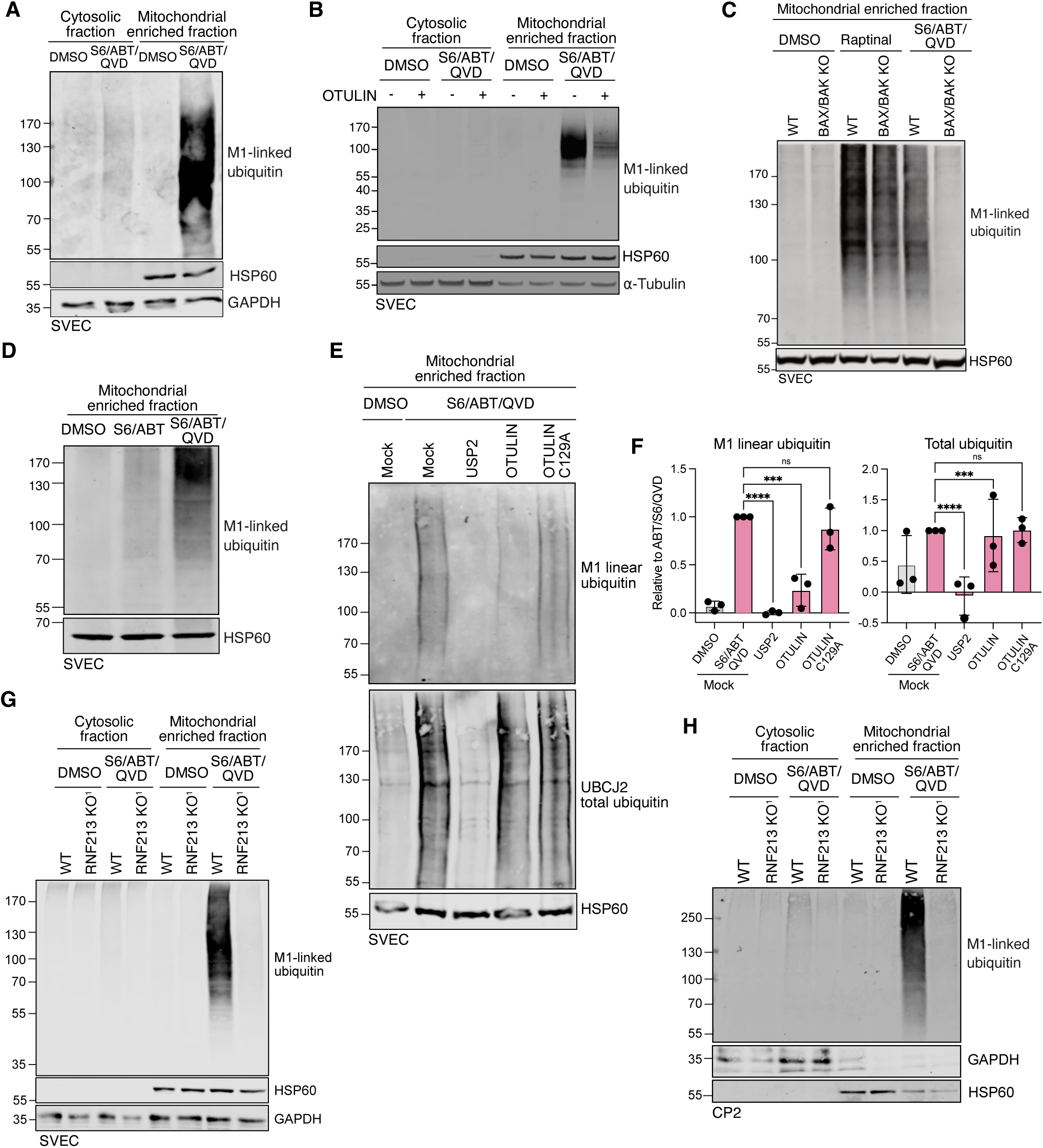
Mitochondrial M1-linked ubiquitination requires RNF213. **A)** SVEC cells were treated +/- ABT-737, S63845 and Q-VD-OPh for 1 hr. Mitochondrial enriched fractions were obtained by digitonin fractionation and immunoblotted for M1-linked ubiquitin. **B)** SVEC WT cells or SVEC cells expressing mScarlet-Otulin were treated with S63845, ABT-737 and Q-VD-OPh for 1hr. Cytosolic/mitochondrial enriched fractions were obtained by digitonin fractionation and immunoblotted for M1-linked ubiquitin. **C)** SVEC WT or BAX/BAK KO cells were treated with either Raptinal (3 hrs) or ABT-737, S63845 and Q-VD-OPh (1 hr) alongside a DMSO control. Mitochondrial enriched fractions were obtained by digitonin fractionation and immunoblotted for M1-linked ubiquitin. **D)** SVEC cells were treated with ABT-737, S63845 and +/- Q-VD-OPh for 1 hr alongside a DMSO control. Mitochondrial enriched fractions were obtained by digitonin fractionation and immunoblotted for M1-linked ubiquitin. **E)** Representative UbiCRest analysis of ubiquitin chains following MOMP. SVEC cells were treated +/- ABT-737, S63845 and Q-VD-OPh for 1 hr. Cells were fractionated by Dounce fractionation and mitochondrial pellets were treated with indicated DUBs or buffer only (Mock) for 1 hr at 37 °C and analyzed for M1-linked and total ubiquitin. **F)** Quantification of M1-linked and total ubiquitin signal remaining after DUB treatment shown relative to Mock S6/ABT/QVD for data represented in A. Graphs display mean values ± SD of n = 3 independent experiments. Statistics were performed using two-way ANOVA **P<0.01, ****P<0.001. **G)** SVEC WT or RNF213 KO^1^ cells were treated +/- ABT-737, S63845 and Q-VD-OPh for 1 hr. Cytosolic/mitochondrial enriched fractions were obtained by Douncer fractionation and immunoblotted for M1-linked ubiquitin. **H)** CP2 WT or RNF213 KO^1^ cells were treated +/-ABT-737, S63845 and Q-VD-OPh for 1 hr. Cytosolic/mitochondrial enriched fractions were obtained by digitonin fractionation and immunoblotted for M1-linked ubiquitination. Where indicated (A – D, G-H), HSP60, α-tubulin and GAPDH were probed as loading controls.

### RNF213 can directly catalyze M1-linked ubiquitination independently of LUBAC

We next asked whether RNF213 ubiquitin ligase activity is required for M1-linked ubiquitination following MOMP. For this purpose, we expressed WT RNF213 or RNF213 with an inactivating mutation (C4516S) in its C-terminal RZ domain, required for ubiquitin ligase activity in MEF ^22^. Cells were treated to undergo MOMP and immunoblotted for M1-ubiquitination (**Fig. 5A**). Extensive M1-linked ubiquitination was detected specifically following MOMP in cells containing WT RNF213, but not the C4516S mutant, demonstrating that the ubiquitination is dependent on RNF213 ligase activity. We next investigated whether RNF213 RZ ubiquitin ligase activity was required for NEMO recruitment following MOMP. Cells co-expressing GFP-fused WT or C4516S RNF213 and mScarlet-NEMO were generated, treated to undergo MOMP and analysed by Airyscan confocal microscopy (**Fig 5B**). Extensive association of mScarlet-NEMO and GFP-RNF213 was specifically observed following MOMP (3^rd^ panel). Furthermore, this association is dependent on RNF213 ubiquitin ligase activity as NEMO recruitment was not observed in the C4516S mutants.

**Figure 5.**
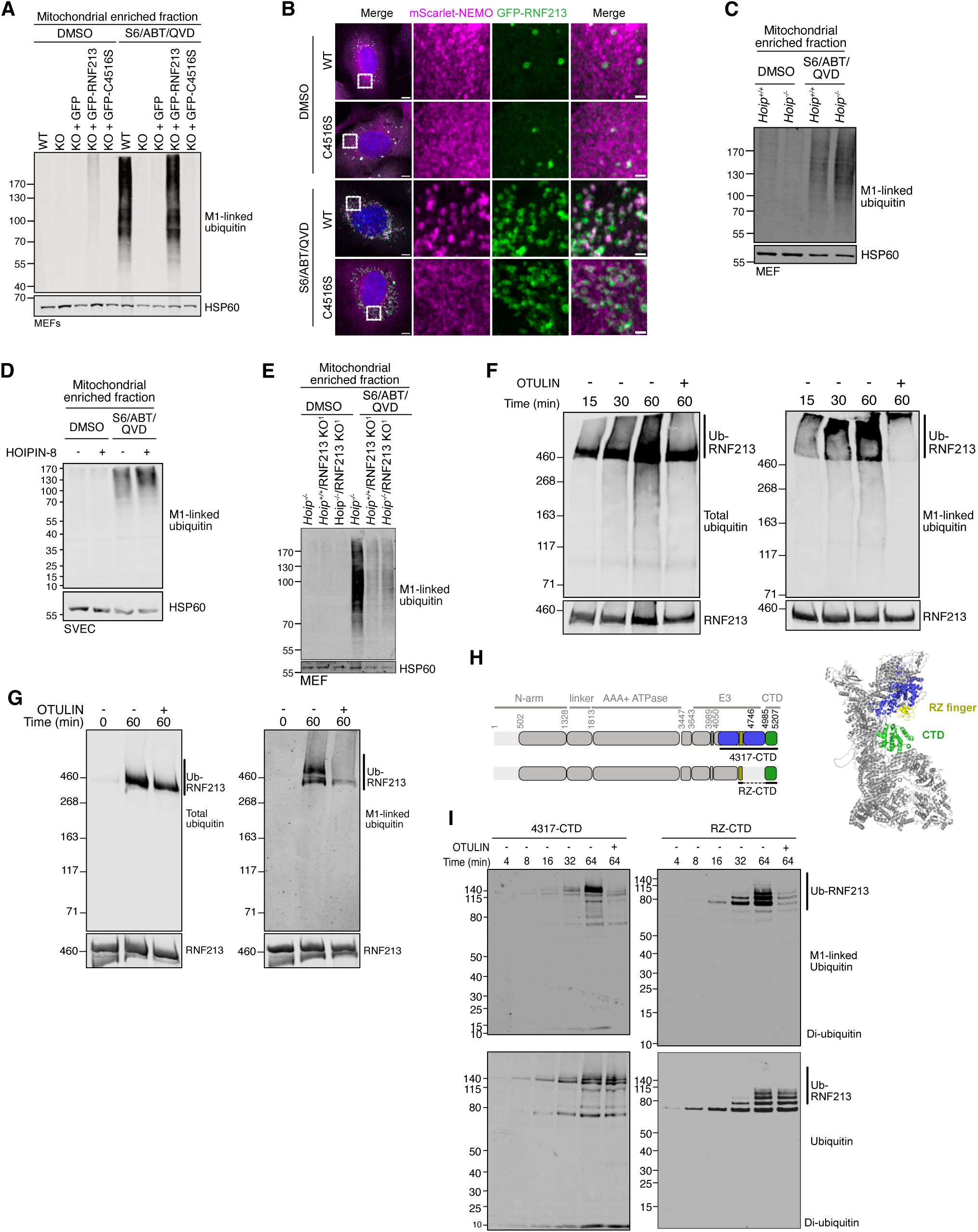
RNF213 can directly catalyse M1-linked ubiquitination independently from LUBAC. **A)** WT, RNF213 KO MEF, and RNF213 KO MEF expressing either dox-inducible GFP, GFP-RNF213 or GFP-C5146S were treated +/- ABT-737, S63845 and Q-VD-OPh for 3 hrs. Mitochondrial enriched fractions were obtained by digitonin fractionation and immunoblotted for M1-linked ubiquitin. **B)** RNF213 KO MEF expressing dox-inducible GFP-RNF213 or GFP-C5146S were treated +/- ABT-737, S63845 and Q-VD-OPh for 3 hrs and imaged by LSM800 confocal microscopy. GFP-NEMO (green), TOM20 (magenta). Brightness and contrast of images were independently adjusted to ensure optimal visualization of protein expression and localisation. Scale bar = 10 µm, insets = 1 µm. **C)** MEF *Tnf*^-/-^/*Hoip^+/+^*or MEF *Tnf*^-/-^/*Hoip^-/-^* cells treated +/- ABT-737, S63845 and Q-VD-OPh for 3 hrs. Mitochondrial enriched fractions were obtained by digitonin fractionation and immunoblotted for M1-linked ubiquitin. **D)** SVEC cells were treated +/- ABT-737, S63845 and Q-VD-OPh for 1 hr +/- LUBAC inhibitor HOIPIN-8. Mitochondrial enriched fractions were obtained by digitonin fractionation and immunoblotted for M1-linked ubiquitin. Where indicated HSP60 (mitochondrial) and α-tubulin were probed as loading controls. **E)** MEF *Tnf*^-/-^/*Hoip^-/-^*and MEF RNF213 KO^1^ *Tnf*^-/-^*/Hoip^+/+^*and MEF *Tnf*^-/-^/*Hoip^-/-^* were treated for 3 hr +/- ABT-737, S63845 and Q-VD-OPh. Mitochondrial enriched fractions were obtained by digitonin fractionation and immunoblotted for M1-linked ubiquitin. **F)** Western blots showing autoubiquitination reactions catalyzed by RNF213 in the presence of UBA1, UBE2L3, ubiquitin and Mg^2+^ -ATP over the indicated times, detected with anti-M1-linked ubiquitin (right panel) or anti-ubiquitin (left panel) antibodies. The 60 min time point reaction was subsequently treated with Otulin for 30 min. **G)** Western blots showing autoubiquitination reactions catalyzed by RNF213 as in F) but using ubiquitin with all lysines mutated (K0-ubiquitin). Autoubiquitination was detected with anti-M1-linked ubiquitin (right panel) or anti-ubiquitin (left panel) antibodies. The 60 min time point reaction was subsequently treated with Otulin for 30 min. **H)** Domain structures of RNF213 showing a truncated fragment (blue) consisting of residues 4317-C (4317-CTD) encompassing the catalytic RZ finger (yellow) and CTD (top) and a minimal catalytic fragment consisting of RZ finger fused to the CTD (green) with a 5xGGSG linker (RZ-CTD; bottom). AlphaFold3 cartoon model of RNF213 residues 474-C (colored in grey) with residue 4317-C colored as in the schematic. **I)** Western blots showing autoubiquitination reactions catalyzed by RNF213 4317-CTD (left) or RZ-CTD (right) in the presence of UBA1, UBE2L3, ubiquitin and Mg^2+^ -ATP over the indicated times, detected with anti-M1-linked ubiquitin (top panels) or anti-ubiquitin (bottom panels) antibodies. The 64 min time point reaction was subsequently treated with Otulin for 30 min.

The canonical E3-ligase required for M1-ubiquitination is the LUBAC protein complex, which is comprised of HOIP, HOIL and SHARPIN ^17, 34–36^. Notably, we previously found that mitochondrial recruitment of NEMO following MOMP was unaffected in cells lacking HOIP and TNF ^13^. To examine whether LUBAC is required for MOMP-dependent M1-linked ubiquitination, we used MEF *Hoip*^-/-^ *Tnf*^-/-^ MEF. Surprisingly, mitochondrial M1-linked ubiquitination was not impacted by the loss of LUBAC or TNF (**Fig. 5C**). To examine a role for LUBAC using an alternative approach, we treated SVEC cells to undergo MOMP in the presence of the HOIP inhibitor HOIPIN-8 ^37^. As expected, TNF-induced M1-linked ubiquitination was effectively inhibited following HOIPIN-8 treatment (**Fig. S4A**). Nonetheless, HOIPIN-8 treatment failed to affect MOMP-dependent mitochondrial M1-linked ubiquitination (**Fig. 5D**), mirroring our findings in LUBAC-deficient MEF cells. We performed similar experiments in LUBAC-deficient cells lacking SHARPIN (*Cpdm^-/-^ Tnf^-/-^* MEF) or HOIL (HOIL KO SVEC) (**Fig. S4B** and **C)**. Aligning with earlier data, absence of either SHARPIN or HOIL did not impact M1-ubiquitination. Given this, we investigated whether RNF213 was required for mitochondrial M1-linked ubiquitination in the absence of LUBAC. *Hoip^-/-^ Tnf*^-/-^ MEF cells lacking RNF213 were generated by CRISPR/Cas9 engineering (**Fig. S4D**). Following engagement of MOMP, M1-linked ubiquitination of mitochondria occurred to the same extent in WT and LUBAC-deficient MEFs – consistent with our previous data. However, RNF213 KO completely prevented mitochondrial M1-linked ubiquitination in *Hoip^-^*^/-^ *Tnf*^-/-^ MEF (**Fig. 5E**).I RNF213 is therefore able to M1-ubiquitinate permeabilized mitochondria following MOMP, independent of the canonical M1-ubiquitnating ligase, LUBAC.

We next addressed whether RNF213 could directly catalyse M1-ubiquitination, using auto-ubiquitination as a proxy assay for its activity. For this purpose, we generated recombinant RNF213 lacking its N-terminal intrinsically disordered region (RNF213 Δ371 ^25^) (**Fig S4E, F**). RNF213 Δ371 was catalytically active, supporting RNF213 autoubiquitination when incubated with E1, the E2 UBE2L3, and ubiquitin (**Fig. 5F**). An increase in M1-linked ubiquitin was detected in the auto-ubiquitinated species in a manner that could be readily removed by treatment with OTULIN, indicating that RNF213 can catalyze the formation of M1-linked ubiquitin chains (**Fig. 5F**). OTULIN treatment did not remove the majority of autoubiquitination detected with anti-ubiquitin antibody, indicating that RNF213 also generates other types of ubiquitin linkages (**Fig. 5F**). To further test the ability of RNF213 to mediate M1-ubiquitination, we performed similar assays using untagged WT and lysine-less (K0) ubiquitin where, in the case of K0-ubiquitin, poly-ubiquitin chain linkages could only occur via linear M1-ubiquitination (**Fig. 5G**). In line with the ability of RNF213 to mediate M1-ubiquitination, auto-ubiquitinated K0-ubiquitin chains were detected by the M1-antibody, and were lost with OTULIN treatment (**Fig 5G)**. Inspection of an AlphaFold model of the structure of human RNF213 shows that the catalytic module encompassing the RZ domain to the C-terminus is connected to a long scaffold and AAA+ ATPase domain (**Fig. 5H**). We assessed whether the catalytic module could be expressed on its own and successfully produced a truncated fragment encompassing the region from just upstream of the RZ finger domain to the C-terminus (residues 4317-C) (**Fig. 5I**, **Fig. S4G**). This fragment (blue) includes the catalytic RZ domain (yellow) and the C-terminal domain (CTD, green) involved in binding E2 conjugated ubiquitin. Notably, this fragment was catalytically active, supporting both autoubiquitination and the formation of free di-ubiquitin chains in the presence of E1, the E2 UBE2L3, and ubiquitin. When the reaction products were probed with an M1-linkage specific antibody, an increase in M1-linked ubiquitin was detected in both the auto-ubiquitinated species and di-ubiquitin products increasing over time (**Fig. 5I**, upper left panel). These products were readily removed by OTULIN, further indicating that RNF213 is capable of catalyzing the formation of M1-linked ubiquitin chains. However, OTULIN addition did not remove the majority of autoubiquitination and di-ubiquitin products, again demonstrating that RNF213 also generates other types of ubiquitin linkages (**Fig. 5I**, **lower left panel**). To further dissect the catalytic requirements for ubiquitin chain formation, we engineered a minimal construct comprising the RZ domain linked to the CTD via a flexible 5xGSGG linker (**Fig. 5I right panels, Fig. S5G**). This minimal fragment retained catalytic activity and could also produce M1-linked ubiquitin chains. Combined, these data show that RNF213 catalyzes M1-linked ubiquitination following MOMP independently of LUBAC, thus enabling NEMO activation and leading to inflammation.

## Discussion

We report that the E3 ubiquitin ligase RNF213 promotes inflammation in response to mitochondrial damage. Following mitochondrial outer membrane permeabilization (MOMP), unexpectedly, RNF213 drives M1-linked ubiquitination, leading to NEMO recruitment and activation of pro-inflammatory NF-κB signalling, thereby directly linking mitochondrial outer membrane integrity with engagement of inflammation.

Loss of mitochondrial membrane potential leading to PINK1 stabilization can also stimulate NF-κB signalling via the E3 ligase Parkin ^38^. We previously excluded a role for the PINK1–Parkin pathway in MOMP-induced inflammation, prompting our search for the relevant E3 ligase machinery ^13^. Combined with our data, this demonstrates that loss of mitochondrial integrity (RNF213) and loss of mitochondrial function (Parkin) engage distinct signalling pathways that converge on NF-κB activation. Beyond mitochondrial damage, RNF213 promotes pro-inflammatory NF-κB signalling in other contexts, including lipotoxic and hypoxic stress, through alternate means ^26, 27^. For instance, during hypoxia, RNF213 activates NF-κB by promoting the degradation of the inhibitory proteins CYLD and SPATA2^27^. Collectively, these findings support a broad role for RNF213 as a regulator of inflammatory signalling.

The specific mechanism of RNF213 activation remains poorly understood. Elucidating this will help provide testable insight into how MOMP activates RNF213. RNF213 is a large protein, with the most common isoform containing two adjacent AAA+ ATPase domains, a RING domain and, identified more recently, a CBM20 carbohydrate-binding domain ^39^. ATP has been proposed as a pathogen activated molecular pattern (PAMP), promoting activation via the AAA+ ATPase domain of RNF213 during infection ^40^. However, MOMP is associated with a reduction in ATP synthesis, therefore making it unlikely that increased ATP is connecting mitochondrial damage with RNF213 activation in this context ^41^. Indeed, reduction of ATP levels as cells die may serve to help inhibit RNF213 activity, limiting inflammation. Consistent with this, we find that whilst RNF213-dependent M1-ubiqutination is detectable during apoptosis, this is significantly enhanced by caspase inhibition.

RNF213 is activated by phylogenetically diverse pathogens including bacteria, viruses and parasites ^20–23^. Nonetheless, how RNF213 recognizes diverse pathogens as well as permeabilized mitochondria remains an outstanding question - conserved damage activated molecular patterns (DAMPs) are lacking. Our data shows that MOMP, either via activated BAX and BAK or by the small-molecule rapitinal, is sufficient to activate RNF213 dependent ubiquitination. Given the ancient bacterial ancestry of the mitochondrial inner membrane, potentially its cytosolic exposure upon outer membrane permeabilization may serve as a DAMP to activate RNF213 and initiate downstream inflammatory signaling.

LUBAC, comprising SHARPIN, HOIL and HOIP, is the canonical E3 ligase complex responsible for M1-linked ubiquitination and is best characterised in TNF signalling ^42^. Consistent with this, we found that TNF-induced M1-linked ubiquitination required LUBAC. Unexpectedly, however, M1-linked ubiquitination following mitochondrial outer membrane permeabilization (MOMP) was LUBAC-independent and instead required RNF213. This resolves the apparent paradox from our earlier observation that NEMO recruitment to mitochondria depends on its binding to M1-linked ubiquitin yet is independent of both LUBAC and TNF ^13^. We find RNF213 ubiquitin ligase activity is essential for MOMP-induced M1-linked ubiquitination, since disruption of its ligase activity through mutation of its C-terminal RZ domain completely abolished M1-linked ubiquitination. Moreover, RNF213 was able to directly catalyse M1-linked ubiquitination *in vitro*.

During *Salmonella* infection, RNF213-mediated ubiquitination of LPS recruits and activates LUBAC, which is then required for M1-linked ubiquitination ^22^. Our data show that RNF213 itself generates a small proportion of total M1-linked ubiquitin relative to total ubiquitin-linkages, both *in vitro* and in cells. Potentially, differences in the magnitude or substrate specificity of RNF213 activity following MOMP versus *Salmonella* infection may therefore determine whether LUBAC is required for M1-linked ubiquitination. Notably, LUBAC-independent M1-linked ubiquitination has been reported in specific intracellular pathogen infection models, where RNF213 has been suggested to contribute, although whether this requires its ligase activity remains unclear ^43, 44^.

Together, our findings demonstrate that M1-linked ubiquitination can occur independently of LUBAC and identify RNF213 as an alternative M1-generating E3 ligase. This raises the exciting possibility that additional E3 ligases may catalyse M1-linked ubiquitination in diverse biological contexts. Crucially, we have now identified RNF213 as the previously unidentified E3 ligase which ubiquitinates mitochondria upon MOMP, providing a key mechanistic link to inflammatory signalling which occurs upon mitochondrial damage.

## Acknowledgments

We thank the Beatson Advanced Imaging Resource (BAIR, RRID:SCR_023875) for help with imaging. We thank Henning Walczak and Gianmaria Liccardi for constructive input and for the provision of HOIP and SHARPIN WT and deficient MEFs; Lifeng Pang for the RNF213 expression plasmids; Rune Busk Damgaard for Otulin expression plasmids; Felix Randow for the GFP-RNF213 expressing MEFs; Genentech for providing the anti-M1 antibody; and Catherine Winchester (CRUK Scotland Institute) for the critical review of this manuscript.

## Funding

Novartis Foundation for Medical-Biological Research (RH)

BBSRC BB/Y01068X/1 (SWGT)

Cancer Research UK Discovery Programme [(DRCNPG-Jun22\100011 and A20145 (SWGT)

Beatson Institute Advanced Technology Facilities A17196 (SWGT)

Beatson Institute Advanced Technology Facilities A29800 (SZ)

Cancer Research UK core funding A29256 (DTH, LB, MM)

The Howat Foundation (GVH and BP)

Cancer Research UK 29754 (GVH and BP)

Prostate Cancer UK Career Acceleration Fellowship TLD-CAF22-012 (AA)

## Author contributions

Study conception: RH and ST conceived the study. Scientific investigation and data analysis: RH, HG, LF, MM, CC, CM, BP, AA, DD, PT, HD, LS, JH, EHY, EV, LB, AR-P, AMG, GC, AB, SL. Supervision: ST, LC, VH, SZ, DH. Manuscript draft: RH, HG, LF, CM, ST. All authors critically reviewed the manuscript.

## Materials and Methods

### Cell culture

SVEC cells were purchased from ATCC. MEF *Tnf^−/−^Hoip^+/+^*, MEF *Tnf^−/−^Hoip^−/−^* cell lines have been described before ^45^. CP2 ^30^ and MC-38 were gifted by Thomas Brunner, University of Konstanz. A549 cells were a gift from Daniel Murphy, University of Glasgow. MEF RNF213-GFP cells were a gift from Felix Randow (University of Cambridge). All cell lines were cultured in DMEM high glucose (#21969035), 10% FBS (Gibco #A5256701), 2 mM glutamine (Gibco #25030081) in 5% CO_2_ at 37 °C.

### Antibodies

The following primary antibodies were used in this study: anti-HA (H9658, Sigma-Aldrich, 1:1000), anti-mitochondrial cocktail (45-7799, Thermo Fisher Scientific, 1:000), anti-M1 (ZRB2114, Sigma-Aldrich, WB: 1:1000, IF: 1:500), anti-GAPDH (2118, Cell Signaling Technology), anti-α-tubulin (Merck, T5168, WB: 1:3000), actin (Sigma-Aldrich, A4700, WB: 1:1000), anti-HSP60 (Cell Signaling Technology, 4870. 1:1000), anti-ubiquitin (UBCJ2, Enzo, ENZ-ABS840, WB: 1:100), anti-M1 Fab fragment, 1F11/3F5/Y102L ^31^ (Genentech, WB 1:1000), anti-Ubiquitin Lys63-specific (Apu3, Merck, 05-1308, WB: 1:100), pIKBα (Cell Signaling Technology, 2859, WB: 1:1000), IKBα (Cell Signaling Technology, 4814, WB: 1:1000), pTBK1 (Cell Signaling Technology, 5483, 1:1000), TBK1 (Cell Signaling Technology, 51872, WB: 1:1000), pSTING (Cell Signaling Technology, 72971, WB: 1:1000), STING (Cell Signaling Technology, 13647, WB: 1:1000), pIRF3 (Cell Signaling Technology, 79945, WB: 1:1000), SMAC (Cell Signaling Technology, 2954, WB: 1:1000) anti-TOM20 (Proteintech, 11802-1-AP, IF, 1:300 or 1:500, Sigma-Aldrich WH0009804M1), anti-RNF213 (Merck, MABN2510, IF 1:100), anti-RNF213 (Sigma-Aldrich, HPA026790, WB 1:1000)

The secondary antibodies used in this study were: goat anti-rabbit IgG (H + L) Alexa Fluor Plus 800 (Invitrogen #A32735), goat anti-mouse IgG (H + L) Alexa Fluor 680 (Invitrogen #A21057), goat anti-mouse IgG (H + L) Dylight 800 (Invitrogen #SA535521), Peroxidase-conjugated AffiniPure donkey anti-human IgG (H+L) (Jackson ImmunoResearch #709-035-149), Alexa Fluor 488 goat anti-rabbit IgG (H + L) (Invitrogen #A11034), Alexa Fluor 488 goat anti-mouse IgG (H + L) (Invitrogen #A11029), Alexa Fluor 568 goat anti-rabbit IgG (H + L) (Invitrogen #A11011), Alexa Fluor 568 goat anti-mouse IgG (H + L) (Invitrogen #A11004), Alexa Fluor 647 goat anti-rabbit IgG (H + L) (Invitrogen #A21245) and Alexa Fluor 647 goat anti-mouse IgG (H + L) (Invitrogen #A21236).

### Chemicals

The following chemicals were used: ABT-737 (APEXBIO, #A8193), S63845 (Chemgood, #C-1370), Q-VD-OPh (AdooQ Bioscience, #A14915-25), mouse recombinant Tnfα (PeproTech, 315-01A), HOIPIN-8 (MedchemExpress, HY-122882), raptinal (Selleckchem, E1560), human IFN-gamma recombinant protein (PeproTech, 300-02).

### Generation of stable cell lines via viral transduction

Over-expression and CRISPR-Cas9 engineered gene knock-out cell lines were generated by retro- and lentiviral infections as described in ^13^. Briefly, HEK293FT cells were transfected using Lipofectamine 3000 according to the manufacturer’s instructions. 1 μg VSVG (Addgene #8454), 1.86 μg psPAX2 (Addgene #12260) and 5 μg plasmid were used for lentiviral infections, 1 μg VSVG and 1.86 μg HIV gag-pol (Addgene #14887) with 5 μg plasmid were used for retroviral infections. After 48 hrs, virus-containing medium was removed from the HEK293FTs and supplemented with 20 μg/ml polybrene before the transfer onto target cells. The selection of infected cells was started 32 hrs later using puromycin, blasticidin or hygromycin at adequate selection concentrations previously tested on parental cells. Alternatively, cells were FACS sorted for the highest 10% GFP- or mScarlet3-expressing cells.

The CRISPR-Cas9 engineered cells were generated using lentiCRISPRv1 or lentiCRISPRv2 vector (Addgene #52961) containing puromycin, blasticidin and hygromycin resistance.

Mouse *RNF213* KO^1^: 5’-GAAGCGGTACATCATACGTG

Mouse *RNF213* KO^2^: 5’-GACCTGGGATCTCTCGTCGA

Mouse *RNF213* KO^3^: 5’-GTGGTGTCATCAACCCCGGA

### Generation of m-Scarlet3-Otulin

The mScarlet3-Otulin fragment was synthesized by GeneArt (Thermo Fisher Scientifc) using the codon-optimized mRNA sequence of the mouse Otulin protein (NCBI identifier: <u>NP_001402977.1</u>) and mScarlet3 ^46^. The fragment was cloned into M6blast (M6blastGFP-NEMO, gifted by Felix Randow) using Not1 and HindIII restriction sites. Viral particles for cell transduction were generated using PMD-OGP and PMD-VSVG plasmid (PMD-OGP and PMD-VSVG, gifted by Felix Randow) and mScarlet3-Otulin in quantities described in the viral transduction section.

### ICE analysis for CRISPR

Genomic DNA was isolated from cells transduced with either the empty vector or CRISPR-Cas9 target sequences using the GeneJET Genomic DNA Purification Kit (Thermo Fisher Scientific, K0721). Using the genomic DNA, the genomic region potentially edited by CRISPR-Cas9, was amplified in a PCR reaction using the Phusion® High-Fidelity DNA polymerase (New England Biolabs, M0530S). The reactions were run on a 1% agarose gel, and bands of the correct size were isolated and purified using the QIAEX II Gel Extraction kit (QIAGEN, 20021). Samples were sequenced by Eurofins genomics and analysed using ICE software by Synthego.

### Primers for genomic DNA amplification

mouse_genomic_*Rnf213*_KO^1^ fw: TGGTGGGGTGGGAAGATCT mouse_genomic_ *Rnf213*_KO^1^ rv: ACGAATTTATTGCAAGTTAAAGGGCAATAACT mouse_genomic_ *Rnf213*_KO^2^ fw: ATGCGAGATGTACTGCACCC mouse_genomic_ *Rnf213*_KO^2^ rv: TCAAACCAGCTTGCAAAGGC mouse_genomic_ *Rnf213*_KO^3^ fw:TGTGAAGTCTTAATGATTTTCAGAAGT mouse_genomic_ *Rnf213*_KO^3^ rv: GAGTGCTGAGGTCACC

### RT-qPCR

RNA isolation was performed as described previously ^13^. Genomic DNA was digested in an additional step by incubating 1-2 ug of RNA with DNAse (Sigma-Aldrich #04716728001) for 15 mins at RT and subsequently heat-inactivating the enzyme at 65 °C for 10 mins. The RNA was reverse transcribed into cDNA using the High-Capacity cDNA Reverse Transcriptase kit (Thermo Fisher Scientific #43-688-13) according to the manufacturer’s instructions using the following thermocyler settings: 25 °C for 10 mins, 37 °C for 120 mins and 85 °C for 5 mins. The RT-qPCR was performed as described previously ^13^.

### cDNA primers

*Mouse Actin 5’-CTAAGGCCAACCGTGAAAAG-3’*

*Mouse Actin 3’-ACCAGAGGCATACAGGGACA-5’*

*Mouse Cxcl1 5’-GGCTGGGATTCACCTCAAGAA-3’*

*Mouse Cxcl1 3’-GAGTGTGGCTATGACTTCGGTT-5’*

*Mouse Mip1a 5’-ACTGCCTGCTGCTTCTCCTACA-3’*

*Mouse Mip1a 3’-ATGACACCTGGCTGGGAGCAAA-5’*

*Mouse Mcp1 5’-CACTCACCTGCTGCTACTCA-3’*

*Mouse Mcp1 5’-GCTTGGTGACAAAAACTACAGC-3’*

*Mouse Csf2 5’-AACCTCCTGGATGACATGCCTG-3’*

*Mouse Csf2 3’-AAATTGCCCCGTAGACCCTGCT-5’*

*Mouse Cxcl10 5’-GGTCTGAGTCCTCGCTCAAG-3’*

*Mouse Cxcl10 3’-GTCGCACCTCCACATAGCTT-5’*

### Multiplex cytokine analysis

Cell supernatants from SVEC WT and SVEC RNF213 KO cells treated for 8 hrs with either DMSO or 10 µM S63845, 10 µM ABT-737 and 30 µM Q-VD-OPh were collected and concentrated 4 times by centrifugation using an Amicon Ultra Centrifugal Filter (Merck, UFC8100) spinning at 800 g for 15 mins. The concentrated supernatant was analyzed by EVE Technologies using a mouse cytokine/chemokine 32-Plex Discovery Assay (MD32). The concentrations of cytokines and chemokines were normalized to the protein concentration, measured by BCA, of each sample.

### Mitochondrial isolation

#### Using digitonin

Pelleted cells were lysed in digitonin lysis buffer (0.25 M sucrose, 700 mM Tris-HCl pH 8 and 100 μg/ml digitonin) for 10 mins on ice. The mitochondrial fraction was pelleted at 3000 g for 5 mins. The supernatant was stored as the cytosolic fraction, and the pellet containing the mitochondrial fraction was resuspended in RIPA lysis buffer and stored on ice for 20 mins before centrifugation for 10 mins at maximum speed (15,000 rpm). The supernatant was taken as a mitochondrial fraction.

#### Using a Dounce homogenizer

Cells were resuspended in mitochondrial isolation buffer (200 mM mannitol, 70 mM sucrose, 10 mM HEPES, 1 mM EGTA, pH 7.0, cOmplete protease inhibitor). After resuspension, cells were homogenized using the Dounce tissue grinder by performing 70 strokes up/down manually and centrifuged at 2000 rpm for 5 mins. The supernatant was collected as the cytosolic fraction, and the pellet was resuspended in mitochondrial isolation buffer and spun down as previously described. The pellet was resuspended in RIPA buffer and placed on ice for 20 mins followed by centrifugation at maximum speed (15,000 rpm) for 10 mins. The supernatant was kept as a mitochondrial fraction.

### OTULIN mitochondrial assay

Mitochondria were isolated from SVEC cells using a Dounce homogenizer as previously described. Isolated mitochondria were resuspended in DUB reaction buffer [50 mM Tris-HCl pH 7.5, 50 mM NaCl, 5 mM DTT] and incubated with 2 µM of either USP2, OTULIN, or OTULIN C129A deubiquitylases for 1 hr to remove either total or M1-linked ubiquitin.

### Western blotting

Cells not subjected to fractionation were scraped and collected by centrifugation at 300 g for 5 mins. The cells were lysed in RIPA buffer (10 mM Tris-HCl (pH 7.4), 150 mM NaCl, 1.2 mM EDTA, 1% Triton X-100 and 0.1% SDS supplemented with cOmplete protease inhibitors and phosphatase inhibitors) for 10-20 mins and proteins were isolated by maximal centrifugation (15,000 rpm) for 10 mins at 4 °C. Protein concentrations were measured (Pierce BCA Protein Assay Kit, Thermo Scientific) and samples prepared using NuPAGE™ LDS Sample Buffer (4X), boiled at 95 °C for 5 mins and loaded onto 8, 10, 12% gels, NuPAGE™ 4–12% Bis-Tris gels or NuPAGE™ 3 to 8%, Tris-Acetate when blotting for RNF213. The SDS PAGE gels were then transferred onto nitrocellulose membranes, which were blocked with 5% milk or BSA in TBS for 1 hr followed by overnight incubation of 1:1000 dilution of primary antibodies in 5% milk of BSA in TBS-T. Membranes were then washed and incubated with secondary antibodies at 1:10,000 dilutions for 1 hr and imaged using an Odyssey CLx Imager.

### Generation of recombinant proteins

RNF213 fragments 4317-CTD (RNF213 residues 4317-C) and RZ-CTD (RNF213 residues 4484-4554 and 4983-5207 connected by 5x GGSG linker) were cloned into pGEX4T1 vector with an N-terminal His-GST tag and TEV cleavage site. Both fragments were produced in *E. coli* BL21(DE3) Gold cells (Agilent Cat #230132). Bacteria were grown at 37 °C until an OD_600_ of 0.7-1.0 before expression was induced with 0.2 mM IPTG and continued overnight at 19 °C. Cells were resuspended and lysed in 25 mM Tris-HCl (pH 7.6), 200 mM NaCl, 20 mM imidazole, 5 mM β-mercaptoethanol (BME) and 2.5 mM phenylmethylsulfonyl fluoride (PMSF). Clarified lysates were passed over 10 ml high density nickel resin (Agarose Bead Technologies, 6BCL-QHNi), washed in 25 mM Tris-HCl (pH 7.6), 200 mM NaCl, 20 mM imidazole, 5 mM BME, and eluted in 25 mM Tris-HCl (pH 7.6), 200 mM NaCl, 200 mM imidazole, 5 mM BME. The eluted products were further purified on 10 ml glutathione agarose resin (Agarose Bead Technologies, 4B-GLU), and finally a 120 ml HiLoad 16/600 Superdex 200pg gel filtration column (Cytiva) into buffer containing 25 mM Tris-HCl (pH 7.6), 200 mM NaCl, 1 mM dithiothreitol (DTT).

The near full-length RNF213 mammalian expression construct, pCAGGS-3xFlag-6xHis-3C-RNF213 (371-5207), was provided by Prof. Lifeng Pan and was purified as described in their study (Zhou *et al*, 2025). Briefly, HEK293F cells were grown in FreeStyle 293 expression medium (Gibco, Life Technologies, 12338-018) in a shaker incubator at 37 °C, 120 rpm, 8% CO_2_ and 70% humidity. 1 mg plasmid DNA was transfected into 1L HEK293F cells for 4 days using FreeStyle MAX reagent (Gibco, ThermoFisher Scientific, 16447100). The transfected cells were collected, lysed and spun at 48,000 g for 1 h and the supernatant incubated with anti-FLAG beads (GenScript, L00432) for 2 h at 4 °C. The beads were washed in buffer (50 mM HEPES (pH 7.5), 200 mM KCl, 1 mM DTT) and then incubated overnight with HRV 3C protease at 4 °C to remove the 3xFLAG-6xHis tag. The flow-through and wash from the beads were concentrated using an Amicon Ultra 15 ml 100 kDa cut-off centrifugal concentrator (Millipore, UFC910024). RNF213 was then further purified over a Superose 6 increase 10/300 GL gel filtration column (Cytiva, 29091596).

The pGEX-6P-1-OTULIN(08-348)-C129A construct was a kind gift from Rune Busk Damgaard ^47^. IMAC purification was carried out according to their guidelines, with protein purified using a HisTrap^TM^ FF 5ml column. After gel filtration, clean fractions were pooled into the final stock of purified protein which was then used in future assays.

Tagless wild type ubiquitin (residue 1-76) was cloned into Novagen® pET-3a vector (MilliporeSigma/Sigma-Aldrich 69742) and transformed into Novagen® Rosetta2(DE3)pLysS cells (MilliporeSigma/Sigma-Aldrich 71403). Bacteria were grown at 37 °C until an OD_600_ of 0.7-1.0 before expression was induced with 0.2 mM IPTG and continued overnight at 25 °C. Cells were resuspended and lysed in 50 mM Tris-HCl (pH 7.6), 15 mM MgCl_2_, 0.02% Triton-X 100, and 2.5 mM PMSF. Clarified lysate (50 ml) was subjected to drop wise addition of 70% perchloric acid (0.5 ml) with stirring on ice, followed by centrifugation to remove the precipitate. Next, the supernatant was dialysed against 50 mM Na acetate (pH 4.5, adjusted with glacial acetic acid) at 4 °C, passed through 0.45 μm filter, and purified on a 5 ml HiTrap SP HP (Cytiva) column using 50 mM Na acetate (pH 4.5) with a NaCl gradient from 0-0.4 M. Fractions containing ubiquitin were pooled, adjusted to pH ∼7.0 with 1M Tris-HCl (pH 7.6), concentrated, and purified on a 120 ml HiLoad 16/600 Superdex 200pg gel filtration column in buffer containing 50 mM HEPES (pH 7.5), 150 mM NaCl.

K0-ubiquitin (all lysines mutated to arginines) with an intact N-terminal methionine was made by expressing 6xHis-SUMO-ubiquitin-K0 (pET28) in 6 L of BL21 *E. coli* at 37 °C and expression was induced with 0.2 mM IPTG and continued overnight at 20 °C. Cells were lysed and clear lysates passed over a 5 ml HisTrap FF column (Cytiva, 17525501) and the eluted protein was cut with SUMO protease overnight at 4 °C to remove His-SUMO tags. Undigested His-SUMO-ubiquitin was removed by passing the protein back over the HisTrap FF column. The flow through containing pure K0-ubiquitin was concentrated.

UBE2L3 was cloned into pGEX4T1 vector with an N-terminal GST tag followed by a TEV cleavage site. The resulting construct was transformed into *E. coli* BL21(DE3) Gold cells (Agilent Cat #230132). Cells were grown at 37 °C until an OD_600_ of 0.7-1.0 before expression was induced with 0.2 mM IPTG and continued overnight at 20 °C. Cells were resuspended and lysed in 50 mM Tris-HCl (pH 7.6), 200 mM NaCl, 1 mM DTT and 2.5 mM PMSF. Clarified lysate was applied on glutathione agarose resin and washed with 50 mM Tris-HCl (pH 7.6), 200 mM NaCl, 1 mM DTT. UBE2L3 was released from the resin by cleavage with TEV protease and subsequently purified on a HiLoad 16/600 Superdex 75pg gel filtration column in buffer containing 25 mM Tris-HCl (pH 7.6), 150 mM NaCl and 1 mM DTT.

UBA1, K48-linked di-ubiquitin and K63-linked di-ubiquitin were purified as described previously ^48^. K11-linked di-ubiquitin was obtained using the same protocol as described in ^48^ but with the E2 UBE2S. Linear di-ubiquitin was cloned into Novagen® pRSFDuet™-1 vector with an N-terminal His tag followed by a TEV cleavage site. The resulting construct was transformed into *E. coli* BL21(DE3) Gold cells (Agilent Cat #230132). Cells were grown at 37 °C until an OD_600_ of 0.7-1.0 before expression was induced with 0.2 mM IPTG and continued overnight at 20 °C. Cells were resuspended and lysed in PBS, 330 mM NaCl, 20 mM imidazole, 5 mM BME and 2.5 mM PMSF. Clarified lysate was subjected to purification by nickel-affinity chromatography (high density nickel resin, Agarose Bead Technologies, 6BCL-QHNi), followed by HiLoad 16/600 Superdex 75pg gel filtration column into PBS.

Protein concentration was determined by BioRAD protein assay or by using molar extinction coefficient at 280 nm (ε_280_ = 1,490 cm^-1^*M^-1^ for Ub, ε_280_ = 4,470 cm^-1^*M^-1^ for linear di-ubiquitin).

### RNF213 ubiquitination assay

RNF213 autoubiquitination assays were performed in the presence of 0.2 µM UBA1, 5 µM UBE2L3, 300 µM ubiquitin and 5 µM RNF213 in a buffer containing 50 mM Tris-HCl (pH 7.5), 50 mM NaCl, 5 mM MgCl_2_, 5 mM ATP at RT. At various time points, 6 µl of the reaction was removed and stopped with LDS sample buffer. One 64-min of each reaction was also treated with 10 µM Otulin M1 Deubiquitinase at 37 °C for 30 mins. Samples were run on NuPAGE 4-12% Bis-Tris SDS-PAGE (Invitrogen) in MOPS buffer and blotted to Nitrocellulose membrane (Cytiva) and probed with anti-M1 Linear Ubiquitin (ZRB2114-4X26UL, Sigma-Aldrich) or anti-ubiquitin (sc-8017, Santa Cruz).

RNF213 (371-5207) autoubiquitination assays were performed in the presence of 0.2 µM UBA1, 5 µM UBE2L3, 300 µM ubiquitin (or 100 µM K0-ubiquitin) and 0.35 µM RNF213 in a buffer containing 50 mM Tris-HCl (pH 7.5), 50 mM NaCl, 5 mM MgCl_2_, 5 mM ATP at 37 °C. At various time points, 6 µl of the reaction was removed and stopped with LDS sample buffer. One 60-min sample of each reaction was also treated with 10 µM Otulin M1 Deubiquitinase at 37 °C for 30 mins. Samples were run on NuPAGE 3-8% Tris-Acetate SDS-PAGE (Invitrogen) in Tris-Acetate buffer and blotted to Nitrocellulose membrane (Cytiva) and probed with anti-M1 Linear Ubiquitin (ZRB2114-4X26UL, Sigma-Aldrich) or anti-ubiquitin (sc-8017, Santa Cruz). Blots were re-probed with anti-RNF213 (Sigma-Aldrich HPA026790, 1:1000) to check equal loading.

### Immunofluorescent staining

After treatment cells were fixed using 4% PFA solution for 15 mins, then permeabilized using 0.2% Triton X-100 for 10 min at RT. Coverslips were blocked in 2% BSA in PBS for 1 hr and incubated with primary antibody in 2% BSA in PBS overnight. Coverslips were washed with PBS and incubated for 1 hr with secondary antibody in 2% BSA in PBS. Finally, coverslips were washed with PBS and H_2_O and mounted using ProLong^TM^ Glass Antifade Mountant (Invitrogen #P36980).

### Airyscan and super-resolution microscopy

Fixed cells were imaged using a Zeiss LSM 880 point-scanning confocal microscope on an inverted Zeiss Axio Observer.Z1 with Airyscan detector. Images were acquired sequentially using a 63×/1.4 Plan-Apochromat lens with Zeiss Immersol 518F oil using the 405 nm, 488 nm, 561nm and 633 nm laser lines. Images were acquired sequentially using the optimal resolution (2 x Nyquist) determined by the Zeiss ZEN LSM 2.1 Black software (RRID:SCR_018163) and using channel-specific band-pass emission filters. When acquiring z-stacks, the software-recommended slice interval was used. Airyscan processing was performed using the Airyscan processing function at default settings in the ZEN software, and to maintain clarity some images were pseudocloured and brightness and contrast altered in Fiji (ImageJ v2.0.0). For image analysis Fiji/ImageJ version 2.9.0/1.53t was used ^49^. For the quantification of GFP-NEMO puncta on mitochondria, at least 50 cells/experiment were analyzed manually and scored. Results were plotted using GraphPad Prism 10.

### Lattice Structural illumination microscopy (SIM)

Fixed cells were imaged using an inverted Zeiss Elyra7 widefield microscope equipped with two PCO edge 4.2M sCMOS cameras. Data were collected using a 63 x 1.4 NA oil immersion objective and 13 SIM phases, using track-specific gratings and z-step size as recommended by the acquisition software. Zeiss 518F oil (23C) was used. All tracks were acquired sequentially. HR Diode and DPSS laser lines 405 nm, 488 nm, 561nm and 633 nm were used, with a LBF 405/488/561/647 and a band-pass emission filter 490-560 + LP640. Zen Blue software (version 3.13) was used to acquire and process the data. SIM data processing used “standard” sharpness and Fast Fit mode. To maintain clarity images were pseudocloured, and brightness and contrast altered in Fiji (ImageJ v2.0.0).

### Imaris Rendering

Images acquired using lattice-SIM microscopy on the Zeiss Elyra7 microscope were processed using Imaris 11.0.1. The Surfaces tool was used to identify components, and a Region of Interest (ROI) was specified by manually drawing around cells on three separate slices of the z-stack (one at either end, and one in the centre). Each channel (excluding DAPI) was then created as a surface within this ROI. Minimum intensity and voxel thresholds were assigned for each repeat. All images in a repeat were then processed using these thresholds. In the Imaris Analysis options ‘Overlapped Volume to Surfaces Surfaces’ measurements were selected, alongside ‘Volume’. Total volume of a component in a cell was then calculated by summing all ‘Volume’ measurements, and total surface overlap by summing the ‘Overlapped Volume to Surfaces Surfaces’ measurements. The percentage of overlapped volume to total volume could then be calculated in order to show changes in association between different components in our treatment conditions.

### Incucyte S3 imaging

Cells were seeded into 24-well plates, incubated overnight and live-cell imaging was performed using an Incucyte® S3 (Sartorius), using a 20x/0.45 Plan Fluor objective lens. Images were captured sequentially. GFP-NEMO was captured using 441-481 nm excitation, 503-544 nm bandwidth with 300 ms exposure time, and propidium iodide using 567-607 nm excitation, 622-704 nm bandwidth with 400 ms exposure time. Brightfield was captured with autoexposure. Images were captured with an image size of 1024 x 1024 pixels, yielding a pixel size of 0.62 x 0.62 µm. 4 positions per well were captured at each timepoint. The Incucyte is located in an incubator (HERAcell 240i, ThermoFisher Scientific) to maintain the cells at 37 °C and 5% CO_2_. Images were acquired using the Incucyte software (Sartorius). For cell death assays, the fluorescent intercalating agent, propidium iodide (PI, 10 µg/ml), was added to the cells alongside treatment to visualize cell death. Images were taken every hour. Cell death was measured using the Incucyte software by quantifying propidium iodide positive nuclei normalized to the starting confluency. For visualization of NEMO recruitment to the mitochondria, SVEC cells expressing GFP-NEMO were treated and imaged every 15 mins for 2 hrs, then every hour.

### Immunopurification of mitochondria and mass spectrometry

Mitochondria were enriched using rapid immunopurification method as described previously ^50^ . Briefly, SVEC cells expressing 3XHA-EGFP-OMP25-MITO-TAG (Addgene: 83356) were plated in 10 cm dishes and treated with either DMSO or 10 μΜ ABT-737, 10 μΜ S63845 and 30 μΜ Q-VD-OPh for 1 hr. All subsequent steps were performed at 4 °C. Cells were washed twice with PBS, collected in KPBS (136 mM KCl and 10 mM KH_2_PO_4_, pH 7.25) and centrifuged at 1000 g for 2 mins. Subsequently cell pellets were resuspended in 1 ml KPBS and homogenized using a small clearance 2 ml tissue grinder (Kimble, 885302) with 30-40 strokes. Homogenate was centrifuged at 1000 g for 2 mins and supernatant was incubated with anti-HA beads (Pierce^TM^, Thermo Fisher, #88836) on an end-over-end rotator for 4 mins. Beads were collected on a magnet and washed 3x with KPBS. Subsequently, bound material was lysed in NP40 buffer (50 mM Tris-HCl (pH 7.4), 150 mM NaCl, 1%NP-40 and 5 mM EDTA.) on ice for 20 mins, centrifuged at full speed for 10 mins after that, supernatant collected for mass spectrometry analysis. Beads were treated to enrich and pre-fractionate peptides as described previously (*36*). Briefly, beads were then resuspended in 1% NP40 and left on ice for 20 mins. Supernatant was recovered by centrifugation, and beads were discarded. Mitochondrial proteins recovered into the supernatant were reduced with 10 mM DTT and subsequently alkylated in the dark with 55 mM Iodoacetamide, both reactions were carried out at RT for 1 hr. Alkylated proteins were captured on MagReSyn Hydroxyl magnetic beads (ReSyn Biosciences) following the manufacturer protocol, and double digested in 100 mM ammonium bicarbonate first using Endoproteinase Lys-C (ratio 1:200 enzyme:substrate) for 1 hr, followed by an overnight trypsin digestion (ratio 1:100 enzyme:substrate). Samples were finally desalted using StageTip ^51^.

### MS Analysis

Peptides resulting from all trypsin digestions were separated by nanoscale C18 reverse-phase liquid chromatography using an EASY-nLC II 1200 (Thermo Scientific) coupled to an Orbitrap Fusion Lumos mass spectrometer (Thermo Scientific). Elution was carried out at a flow rate of 300 nl/min using a binary gradient, into a 50 cm fused silica emitter (New Objective) packed in-house with ReproSil-Pur C18-AQ, 1.9 μm resin (Dr Maisch GmbH), for a total run-time duration of 280 mins. Packed emitter was kept at 50 °C by means of a column oven (Sonation) integrated into the nanoelectrospray ion source (Thermo Scientific). Eluting peptides were electrosprayed into the mass spectrometer using a nanoelectrospray ion source. An Active Background Ion Reduction Device (ESI Source Solutions) was used to decrease air contaminants signal level. The Xcalibur 4.2 software (Thermo Scientific) was used for data acquisition. A full scan was acquired at a resolution of 240000 at 200 m/z, over mass range of 375-1500 m/z. Ions were selected during a 3 sec cycle time using the quadrupole, fragmented in the ion routing multipole, and finally analysed in the Ion trap using a maximum injection time of 10 ms or a target value of 3e4 ions. Former target ions selected for MS/MS were dynamically excluded for 30 sec.

### MS Data Analysis

The MS Raw data were processed with MaxQuant software version 1.6.14.0 ^52^ and searched with Andromeda search engine ^53^ , querying UniProt (*Mus musculus* (25198 entries) ^54^. First and main searches were performed with precursor mass tolerances of 20 ppm and 4.5 ppm, respectively, and MS/MS tolerance of 20 ppm. The minimum peptide length was set to six amino acids and specificity for trypsin cleavage was required. Cysteine carbamidomethylation was set as fixed modification, whereas Methionine oxidation and N-terminal acetylation were specified as variable modifications. The peptide, protein, and site false discovery rate (FDR) was set to 1%. All MaxQuant outputs were analysed with Perseus software version 1.6.2.3 ^55^ . Protein abundance was measured using label-free quantification (LFQ) intensities reported in the ProteinGroups.txt file. Only proteins quantified in three out of four replicates in at least one group, were measured according to the label-free quantification algorithm available in MaxQuant ^56^. Missing values were imputed separately for each column, and significantly enriched proteins were selected using a permutation-bas

### Isolation of peptides containing ubiquitin remnants and mass spectrometry

SVEC cells were treated with DMSO or 10 μM S63845, 10 μM ABT-737 and 30 μM Q-VD-OPh for 3 hrs and peptides with ubiquitin remnant motifs were subsequently isolated using a PTMScan Ubiquitin Remnant Motif (K--GG) as described previously ^13^. Peptides were separated using a nanoscale C18 reverse-phase liquid chromatography then analysed by mass spectrometry as previously described ^13^.

### Proteome sample preparation for RNF213 KO confirmation

SVEC and CP2 cells were lysed using 2% SDS in 100 mM Tris-HCl at pH 8.0. Extracted proteins were reduced using 10mM dithiothreitol (DTT) for 30 minutes at 54°C and then alkylated using 55mM iodoacetamide (IAA) at 25°C for 1 hour in the dark. Alkylated proteins were mixed with conditioned magnetic hydroxyl beads (ReSyn Bioscience) and left to aggregate for 30 minutes in 70% Acetonitrile. Samples were transferred to a magnetic rack, and the beads were washed in 70% and 95% Acetonitrile, and supernatant were discarded. Protein beads aggregates were finally resuspended in 100mM ammonium bicarbonate, and trypsin (Promega) was added with an enzyme: substrate ratio of 1:200, following an overnight incubation at 35°C on a shaker at 1000rpm. Samples were finally desalted using StageTip ^51^.

### Mass spectrometry Data Independent Acquisition (DIA)

Peptides resulting from digestion were separated by nanoscale C18 reverse-phase liquid chromatography using an EASY-nLC II 1200 (Thermo Scientific) coupled to an Orbitrap Fusion Lumos mass spectrometer (Thermo Scientific). Elution was carried out at a flow rate of 300 nl/min using a binary gradient, into a 50 cm fused silica emitter (New Objective) packed in-house with ReproSil-Pur C18-AQ, 1.9 μm resin (Dr Maisch GmbH). Packed emitter was kept at 50 °C by means of a column oven (Sonation) integrated into the nanoelectrospray ion source (Thermo Scientific). An Active Background Ion Reduction Device (ABIRD) was used to decrease air contaminants signal level.

Proteome analysis was carried out over a total runtime of 130 minutes in positive ion mode using data-independent acquisition (DIA). A full scan (FT-MS) over mass range of 340–1050 m/z was acquired at 120,000 resolution at 200 m/z, with a target value of 1e6 ions for a maximum injection time of 246 ms. Higher energy collisional dissociation fragmentation spectra were recorded at 30000 resolution at 200 m/z. All precursors were fragmented using 28 consecutive tMSn scan windows with 24 Da width, allowing for a 0.5 m/z overlap, and covering a mass range from 362 to 1010 m/z. All ions were fragmented using normalised collision energy of 28, for a maximum injection time of 54 ms, or a target value of 3e6 ions. Resulting fragmentation spectra were then measured at a resolution of 30,000 at 250 Th. MS data were acquired using the Xcalibur software (Thermo Fisher Scientific).

### DIA data processing

The Proteome Raw data were processed with Spectronaut version 20.5 ^57^ using directDIA™ analysis querying UniProt ^58^ *Mus musculus* (25,662 entries). Enzyme / Cleavage Rule was set to trypsin/P, allowing up to two missed cleavage sites. Methionine oxidation and N-terminal acetylation were specified as variable modifications, whereas Carbamidomethylation of Cysteine was set as fixed modification. Minor peptide grouping was set to “Modified Sequence”, and Major and Minor Group Quantity were set to “Sum peptide quantity” and “Median precursor quantity” respectively. Protein and precursor PEP cutoff was set to 0.05, single hit proteins were excluded from dataset and all other parameter in were left to default values. *0*

### RNA sequencing

WT and RNF213 KO SVEC cells were treated with DMSO or 10 μM S63845, 10 μM ABT-737 and 30 μM Q-VD-OPh for 3 hrs and RNA extracted using GeneJET RNA isolation kit (Thermo Fisher Scientific #K0732) according to the manufacturer’s instructions. Illumina Stranded mRNA: Quality control of all RNA samples was performed (Agilent Tapestation 4200, High Sensitivity RNA screentape), and only those samples showing RIN values >8 were processed. RNA concentrations were determined by Qubit Fluorometer using the Qubit RNA Broad Range assay (both Thermo Fisher), with 500 ng of total RNA used as initial input. Libraries were then prepared using the manufacturer’s standard procedures (Illumina Stranded mRNA), with Illumina RNA Unique Dual Indexes used to index libraries. Post library QC was then performed using High Sensitivity D1000 screentape (Agilent) for library sizing and profiling and quantified using Qubit High Sensitivity DNA assay. Libraries were then pooled equimolar, to a final concentration of 2 nM prior to sequencing. The library pool was then sequenced on an Illumina NextSeq 2000 instrument, at Paired end 75 bp read length and a depth of approximately 25M reads per sample.

### RNA sequence analysis

Raw FASTQ files were first assessed for quality using *fastqc v0.11.8*, followed by adapter trimming and low-quality base removal using *trimgalore v0.4.4*. Post-trimming quality was re-evaluated to ensure data suitability for downstream alignment. Reads were aligned to the mouse reference genome (GRCm39) using the *hisat2 v2.1.0*; and sorted into bam files using *samtools v1.15.1.5*. Gene-level quantification was performed with *featureCounts* function in *Subread v2.0.1*, generating count matrices for downstream expression analysis. The count matrix was then analyzed in *R v4.4.2*. Differential analysis was carried using the *DESeq2 v1.34.0* package, enrichment analysis was carried out using *clusterProfiler v4.16.0* package, and results were plotted using *ggplot2 v3.5.2* package.

### Statistical analysis

Statistical analyses were performed using Prism 10. All data are mean ± standard deviation (SD). ns= not significant, *P<0.05, **P<0.01, ***P<0.001, ****P<0.0001

---

**Figure S1.**
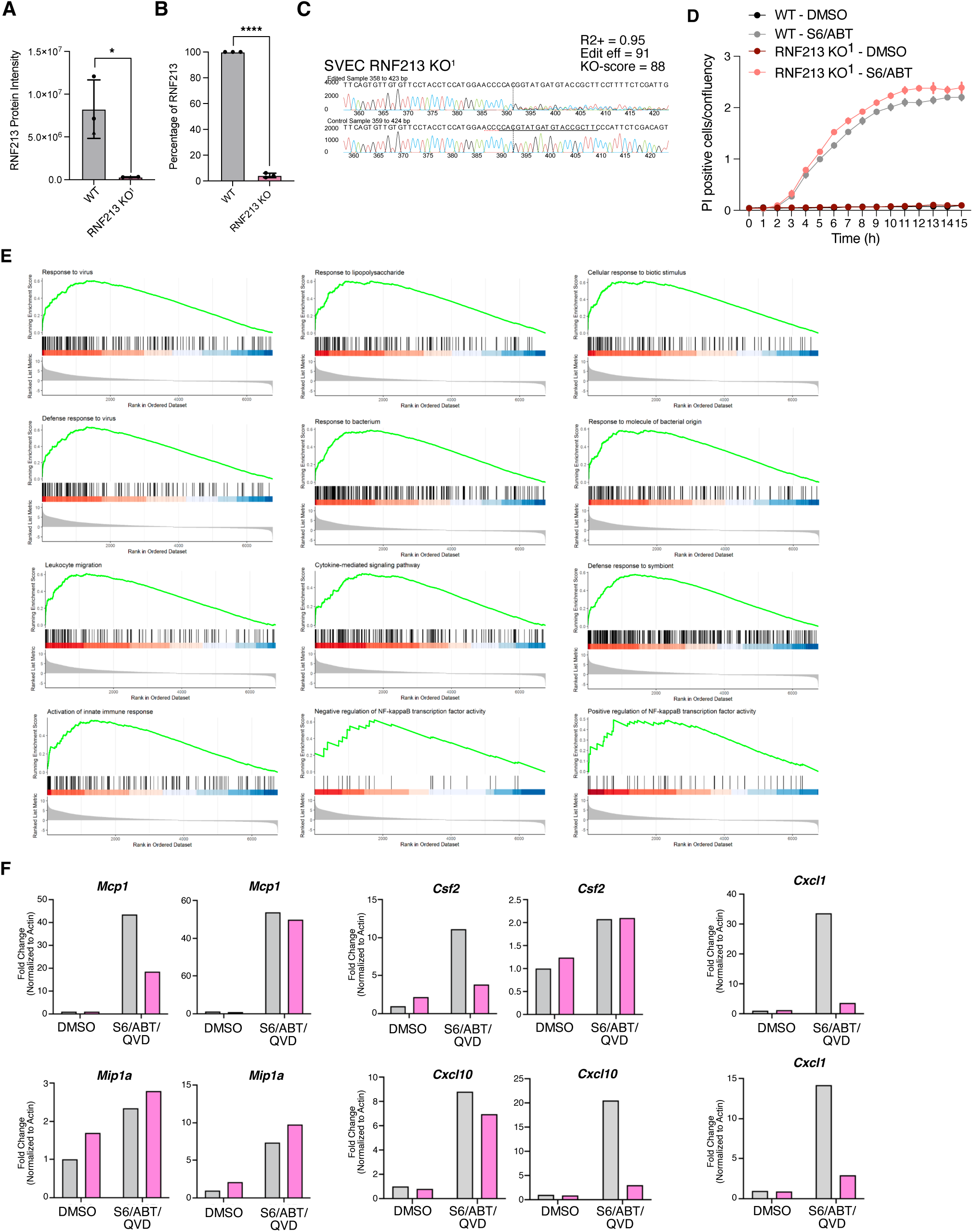
RNF213 is required for NF-κB dependent inflammation following MOMP. **A)** Mass spectrometry was performed on SVEC WT and RNF213 KO^1^ cells. Graph shows the intensity levels of RNF213 protein in each cell line. **B)** Percentage KO of RNF213 in SVEC WT cells compared to RNF213 KO^1^ cells. **C)** Inference of CRISPR Edits (ICE) confirmation of RNF213 knockout via CRISPR in SVEC cells compared to WT control cells. **D)** WT or RNF213 KO^1^ SVEC cells were treated +/- S63845 and ABT-737 then imaged over 15 hr on an S3 using a 20x/0.3NA objective. Propidium iodide (PI) uptake was used to measure cell death, normalized to confluency of cells. Data from one representative experiment shown. **E)** Gene Set Enrichment Analysis (GSEA) of RNAseq analysis carried out on WT and RNF213 KO^1^ SVEC cells treated with S63845/ABT-737/QVD. Pathways associated with inflammatory responses were investigated. RNAseq was carried out on four independent repeats. **F)** SVEC WT or RNF213 KO^1^ cells were treated for 3 hrs +/- ABT-737, S63845 and Q-VD-OPh. Expression of denoted transcripts was measured by qRT-PCR, normalised to actin. Graphs show two of the three independent repeats, complementing the graphs shown in Fig. 2F.

**Figure S2.**
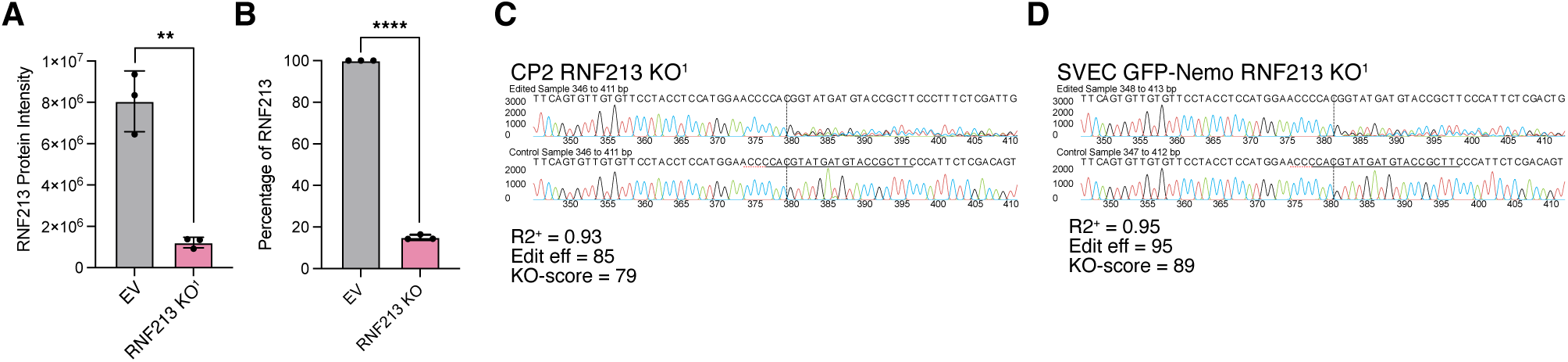
Validation of RNF213 deletion in various cell lines. **A)** Mass spectrometry was performed on CP2 WT and RNF213 KO^1^ cells. Graph shows the intensity levels of RNF213 protein in each cell line. **B)** Percentage KO of RNF213 in CP2 WT cells compared to RNF213 KO^1^ cells. **C)** ICE confirmation of RNF213 knockout (KO^1^) via CRISPR in mouse prostate cancer CP2 cells compared to WT cells. **D)** ICE confirmation of RNF213 knockout (KO^1^) in GFP-NEMO SVEC cells.

**Figure S3.**
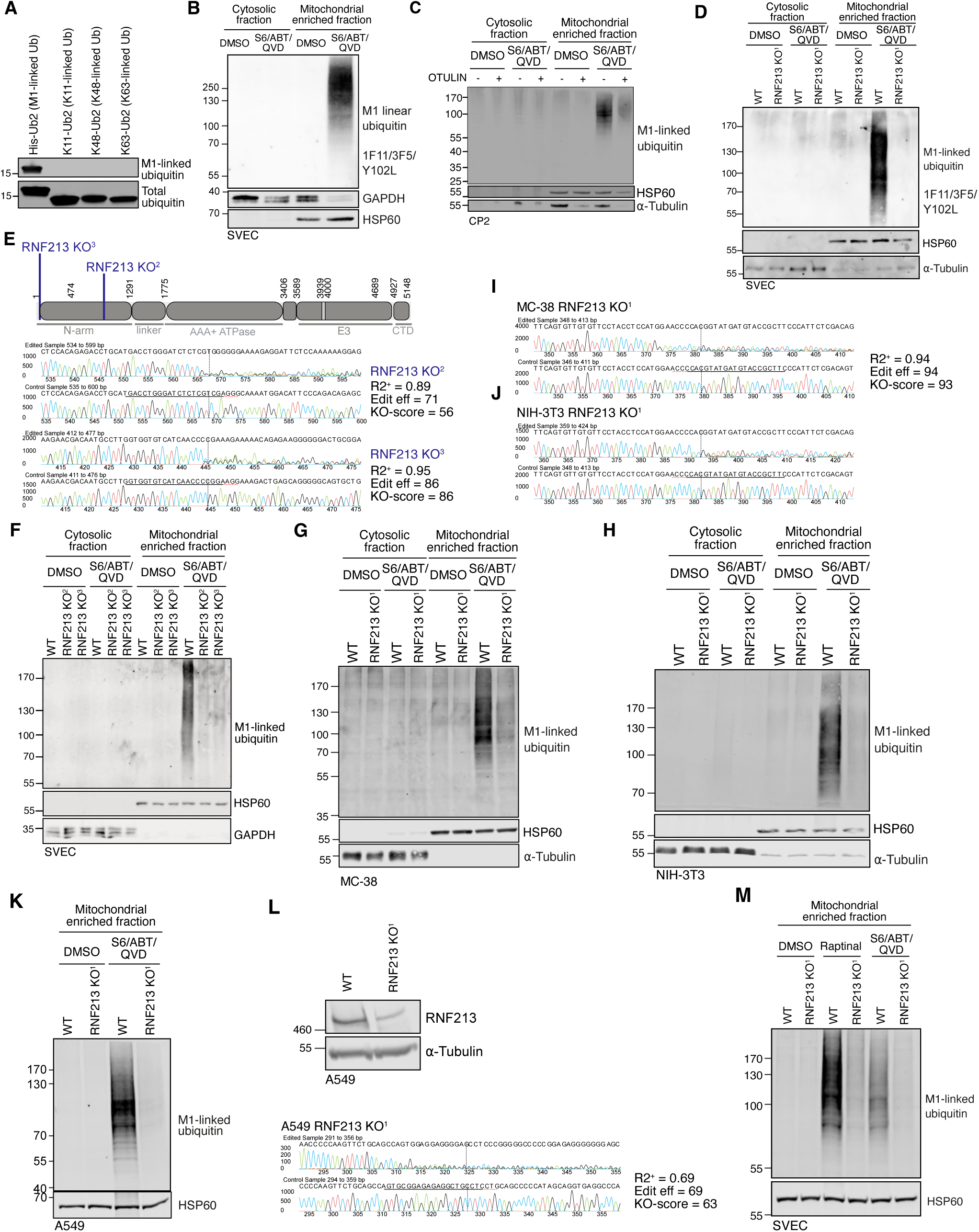
RNF213 promotes M1-linked ubiquitination. **A)** Recombinant di-ubiquitin of different linkages were immunoblotted using the M1 antibody or total ubiquitin. **B)** SVEC cells were treated +/- ABT-737, S63845 and Q-VD-OPh for 1 hr. Cytosolic/mitochondrial enriched fractions were obtained by digitonin fractionation and immunoblotted for M1-linked ubiquitin using an alternative antibody from Genentech (anti-M1 Fab fragment, 1F11/3F5/Y102L). **C)** Control CP2 cells or CP2 cells expressing mScarlet-Otulin were treated +/- ABT-737, S63845 and Q-VD-OPh for 1 hr. Cytosolic/ mitochondrial enriched fractions were obtained by digitonin fractionation and immunoblotted for M1-linked ubiquitin. **D)** SVEC WT or RNF213 KO^1^ cells were treated +/- ABT-737, S63845 and Q-VD-OPh for 1 hr. Cytosolic/mitochondrial enriched fractions were obtained by digitonin fractionation and immunoblotted for M1-linked ubiquitin using an alternative antibody from Genentech (anti-M1 Fab fragment, 1F11/3F5/Y102L). **E)** Schematic of RNF213 protein with guide RNA 2 and 3 used in for CRISPR experiments indicated in blue. ICE confirmation of RNF deletion using additional RNF213 CRISPR knockout guides in SVEC cells compared to WT cells. **F)** SVEC WT and RNF213 KO^2^ or RNF213 KO^3^ cells were treated +/- ABT-737, S63845 and Q-VD-OPh for 1 hr. Cytosolic/mitochondrial enriched fractions were obtained by digitonin fractionation and immunoblotted for M1-linked ubiquitin. **G)** MC-38 WT or RNF213 KO^1^ cells treated +/- ABT-737, S63845 and Q-VD-OPh for 1 hr. Cytosolic/mitochondrial enriched fractions were obtained by digitonin fractionation and immunoblotted for M1-linked ubiquitin. **H)** NIH-3T3 WT or RNF213 KO^1^ cells were treated +/- ABT-737, S63845 and Q-VD-OPh for 3 hrs. Cytosolic/mitochondrial enriched fractions were obtained by digitonin fractionation and immunoblotted for M1-linked ubiquitin. **I)** ICE Validation of RNF213 KO^1^ in MC-38 cells. **J)** ICE Validation of RNF213 KO^1^ in NIH-3T3 cells. **K)** A549 WT or RNF213 KO^1^ cells were IFN stimulated overnight and treated +/- ABT-737, S63845 and Q-VD-OPh for 3 hr. Mitochondrial enriched fractions were obtained by digitonin fractionation and immunoblotted for M1-linked ubiquitin. **L)** A549 WT or RNF213 KO^1^ total cell lysates were immunoblotted for RNF213. ICE Validation of RNF213 KO^1^ in A549 cells. **M)** SVEC WT or RNF213 KO^1^ cells were treated with either Raptinal (3 hrs) or ABT-737, S63845 and Q-VD-OPh for 1 hr alongside a DMSO control. Mitochondrial enriched fractions were obtained by digitonin fractionation and immunoblotted for M1-linked ubiquitin.

**Figure S4.**
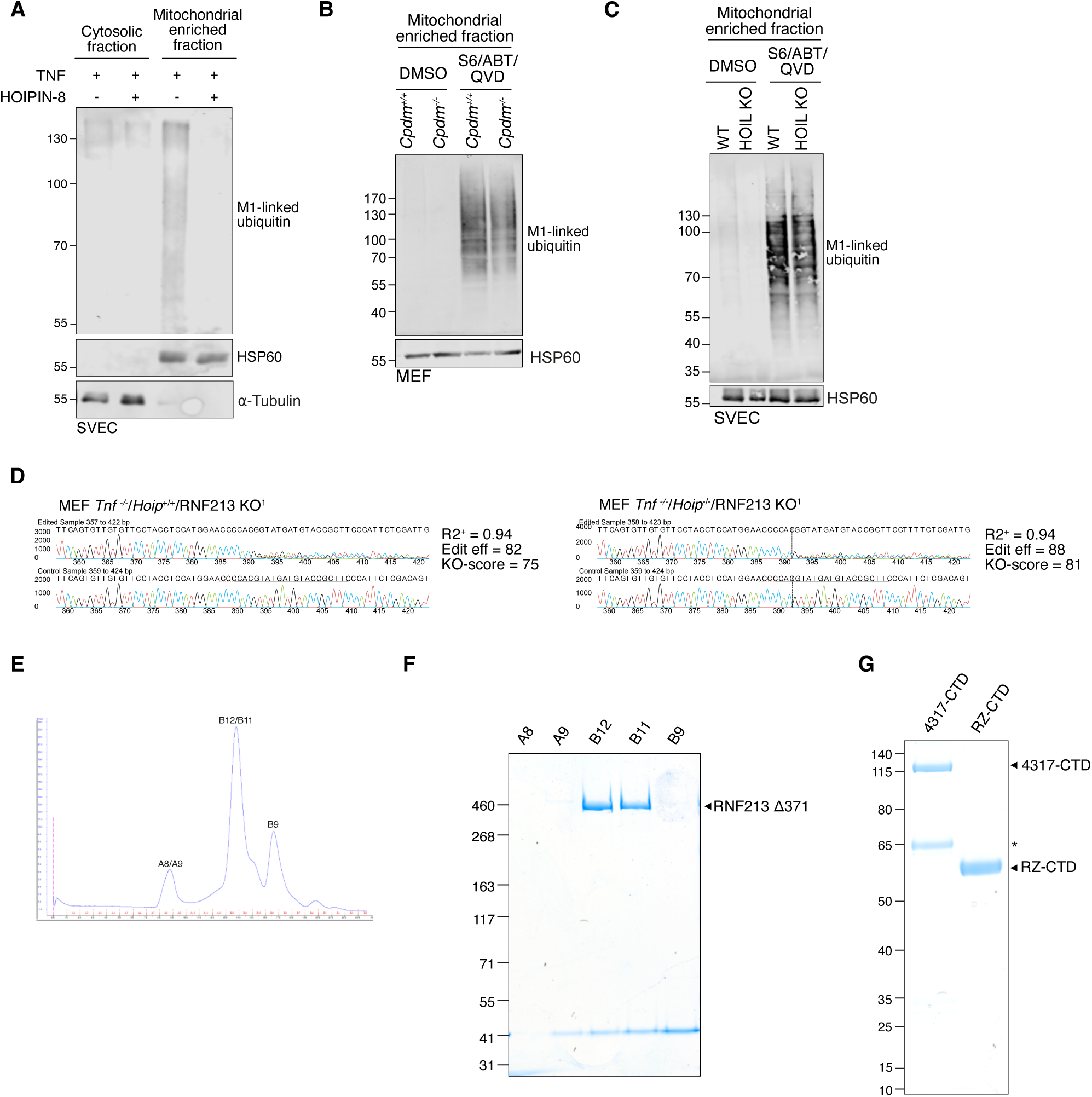
RNF213 catalyzes linear ubiquitination without LUBAC involvement. **A)** SVEC cells treated with 20ng/µl TNFα alongside +/- 30µM HOIPIN-8 for 1 hr Cytosolic/mitochondrial enriched fractions were obtained by digitonin fractionation and immunoblotted for M1-linked ubiquitin. HSP60 (mitochondrial) and α-tubulin were probed as loading and fractionation controls. **B)** MEF *Tnf*^-/-^/*Cpdm^+/+^*or MEF *Tnf*^-/-^/*Cpdm^-/-^* cells were treated +/- ABT-737, S63845 and Q-VD-OPh for 3 hr. Mitochondrial enriched fractions were obtained by digitonin fractionation and immunoblotted for M1-linked ubiquitin. **C)** SVEC WT or HOIL KO cells were treated +/- ABT-737, S63845 and Q-VD-OPh for 1 hr. Mitochondrial enriched fractions were obtained by digitonin fractionation and immunoblotted for M1-linked ubiquitin. **D)** Inference of CRISPR Edits (ICE) confirmation of RNF213 knockout via CRISPR in MEF *Tnf*^-/-^/*Hoip^+/+^* or MEF *Tnf*^-/-^/*Hoip^-/-^*cells compared to WT control cells. **E-G)** RNF213 was expressed *from E. coli* BL21(DE3) E.coli and purified over a 10ml nickel column, followed by a 10ml glutathione column and finally over a 120ml 16/600 Superdex 200pg gel filtration column. **E)** Gel filtration chromatogram of RNF213 purification. **F)** 2µg of each protein fraction was run on a 3-8% Tris-acetate gel in Tris-acetate buffer at 150V for 60 mins, then stained with Coomassie stain. **G)** 2µg of RNF213 protein fragments were run on a 4-12% NuPAGE Bis-Tris gel in MOPS buffer at 200V for 30 mins, then stained with Coomassie stain. The 65kDa band (*) was identified by mass spec. as a proteolytic fragment of RNF213. Experiments were performed for a minimum of n = 3 independent repeats.

